# Human KIR^+^CD8^+^ T cells target pathogenic T cells in Celiac disease and are active in autoimmune diseases and COVID-19

**DOI:** 10.1101/2021.12.23.473930

**Authors:** Jing Li, Maxim Zaslavsky, Yapeng Su, Michael J. Sikora, Vincent van Unen, Asbjørn Christophersen, Shin-Heng Chiou, Liang Chen, Jiefu Li, Xuhuai Ji, Julie Wilhelmy, Alana M. McSween, Brad A. Palanski, Venkata Vamsee Aditya Mallajosyula, Gopal Krishna R. Dhondalay, Kartik Bhamidipati, Joy Pai, Lucas B. Kipp, Jeffrey E. Dunn, Stephen L. Hauser, Jorge R. Oksenberg, Ansuman T. Satpathy, William H. Robinson, Lars M. Steinmetz, Chaitan Khosla, Paul J. Utz, Ludvig M. Sollid, James R. Heath, Nielsen Q. Fernandez-Becker, Kari C. Nadeau, Naresha Saligrama, Mark M. Davis

## Abstract

Previous reports show that Ly49^+^CD8^+^ T cells can suppress autoimmunity in mouse models of autoimmune diseases. Here we find a markedly increased frequency of CD8^+^ T cells expressing inhibitory Killer cell Immunoglobulin like Receptors (KIR), the human equivalent of the Ly49 family, in the blood and inflamed tissues of various autoimmune diseases. Moreover, KIR^+^CD8^+^ T cells can efficiently eliminate pathogenic gliadin-specific CD4^+^ T cells from Celiac disease (CeD) patients’ leukocytes *in vitro*. Furthermore, we observe elevated levels of KIR^+^CD8^+^ T cells, but not CD4^+^ regulatory T cells, in COVID-19 and influenza-infected patients, and this correlates with disease severity and vasculitis in COVID-19. Expanded KIR^+^CD8^+^ T cells from these different diseases display shared phenotypes and similar T cell receptor sequences. These results characterize a regulatory CD8^+^ T cell subset in humans, broadly active in both autoimmune and infectious diseases, which we hypothesize functions to control self-reactive or otherwise pathogenic T cells.

**One-Sentence Summary:** Here we identified KIR^+^CD8^+^ T cells as a regulatory CD8^+^ T cell subset in humans that suppresses self-reactive or otherwise pathogenic CD4^+^ T cells.

---

While most CD8^+^ T cells are geared towards the control of pathogen-infected or cancerous cells, there have been long standing evidence in mice that a small subset can also suppress autoimmune responses (*1*). This regulatory function of CD8^+^ T cells was first implicated by the depletion of CD8^+^ T cells in experimental autoimmune encephalomyelitis (EAE), a mouse model of human Multiple Sclerosis (MS) (*2, 3*). In particular, Cantor and colleagues showed that in B6.Qa-1-D227K mice, disruption of Qa-1-CD8 coreceptor binding leads to spontaneous autoimmune diseases (*4, 5*). The inhibitory C-type lectin-like family of receptors, Ly49, which are ubiquitous on natural killer (NK) cells, were identified as unique surface markers for this regulatory CD8^+^ T cell subset (*6*), and the transcription factor Helios as an essential control element for their differentiation and function in mice (*7*). Recently, our group found that clonally expanded CD8^+^ T cells in EAE recognized peptides bound to H2-D^b^ and that these peptides stimulated Ly49^+^CD8^+^ regulatory T cells and suppressed the disease (*8*). This extends the original observations beyond Qa-1 to encompass classical class I MHC interactions, as does the results of Sheng *et al*. (*9*), suggesting a general mechanism of peripheral tolerance. Here, we identify CD8^+^ T cells expressing inhibitory Killer cell Immunoglobulin like Receptors (KIR), the functional counterpart of the mouse Ly49 family in humans (*10*), as a regulatory CD8^+^ T cell subset in humans that suppresses pathogenic CD4^+^ T cells in Celiac disease (CeD), and likely other autoimmune disorders and infectious diseases as well.

## Increased KIR^+^CD8^+^ T cells in human autoimmune diseases

Both mouse Ly49 and human KIR receptors bind to class I MHC molecules, typically having the inhibitory tyrosine-based inhibition motifs (ITIM) in their cytoplasmic tails, and are ubiquitously expressed on NK cells, as well as a small subset (1∼5%) of CD8^+^ T cells (*6, 10*). Because of these similarities, we analyzed CD8^+^ T cells expressing inhibitory KIRs (hereby designated as KIR^+^CD8^+^ T cells) (*11, 12*) in the peripheral blood of patients with autoimmune diseases and age/gender-matched healthy controls (HC). Since CD3^+^CD56^+^ cells are likely NKT cells which might have a very different antigen recognition mechanism compared to CD8^+^ T cells, we only focused on the conventional CD8^+^ T cells by gating on CD3^+^CD56^-^TCRαβ^+^CD8^+^ cells in this study. KIR3DL1 and KIR2DL3 are the two major KIR subtypes expressed by a small subset of human CD8^+^ T cells (fig. S1). This normally very small subset (1-2%) of KIR^+^CD8^+^ T cells were significantly increased in the blood of many patients with MS, Systemic Lupus Erythematosus (SLE) or CeD, as compared to healthy controls (Fig. 1A).

**Fig. 1.**
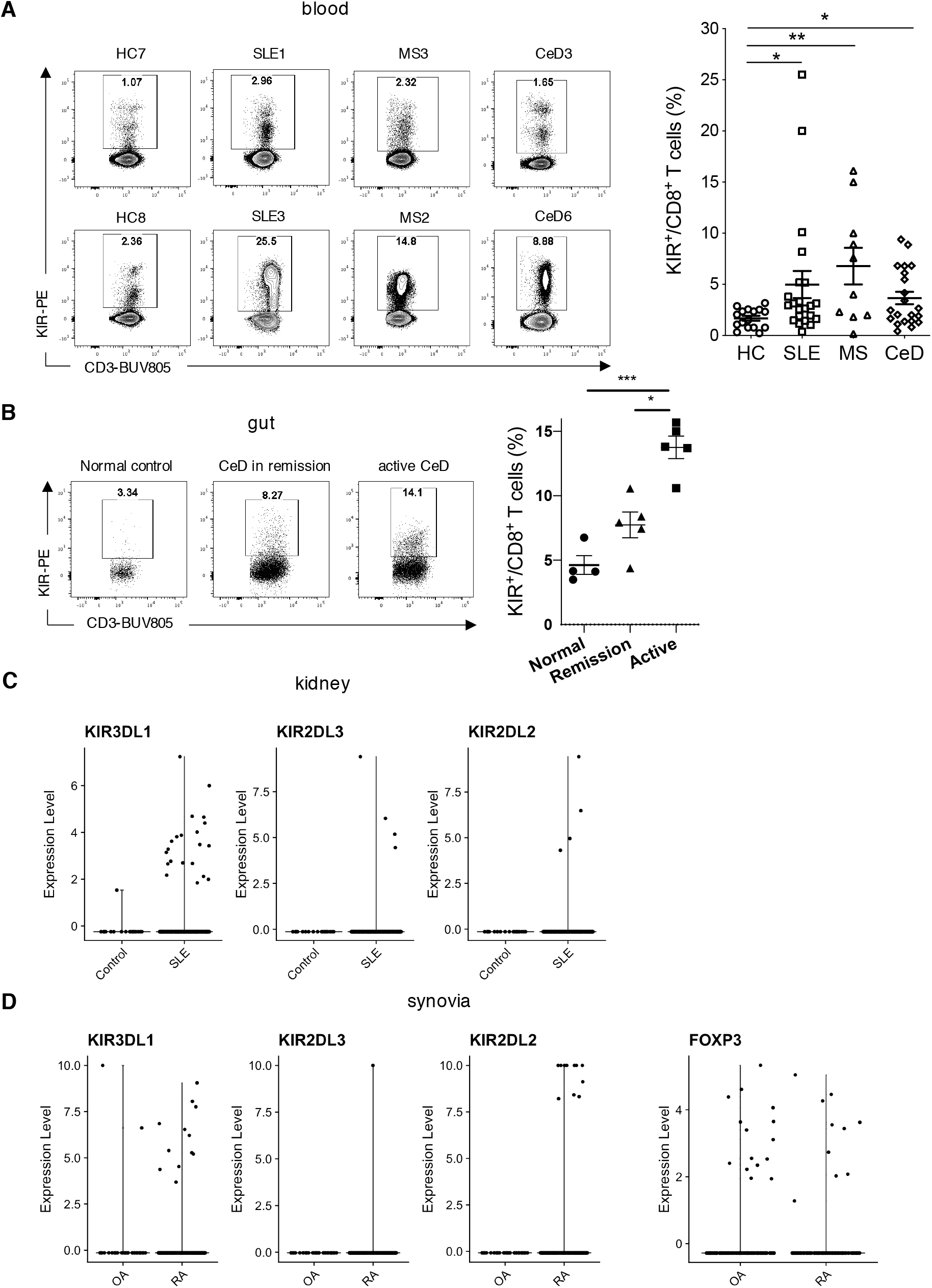
Increased KIR^+^CD8^+^ T cells in patients with autoimmune diseases. (**A**) Representative contour plots (left) and summary histogram (right) showing frequency of KIR^+^ CD8^+^ T cells in the peripheral blood of healthy controls (HC, N=16) and patients with systemic lupus erythematous (SLE, N=22), multiple sclerosis (MS, N=10) or celiac disease (CeD, N=21) analyzed by flow cytometry. KIR^+^ cells were detected by PE-conjugated antibodies against KIR2DL1 (clone#143211), KIR2DL2/L3 (Dx27), KIR2DL5 (UP-R1), KIR3DL1 (Dx9) and KIR3DL2 (clone#539304). \**P*<0.05, \*\**P*<0.01, Kruskal-Wallis test corrected for multiple comparisons. (**B**) Representative contour plots and summary histogram showing frequency of KIR^+^CD8^+^ T cells in the duodenum of normal controls (N=4), CeD in remission (N=5) and active CeD (N=5) analyzed by flow cytometry. \**P*<0.05, \*\*\**P*<0.001, Kruskal-Wallis test corrected for multiple comparisons. (**C**) Expression of *KIR* transcripts (*KIR3DL1, KIR2DL3* and *KIR2DL2*) in CD8^+^ T cells from healthy kidneys (control, N=6) versus SLE nephritis kidneys (N=20) is shown. KIR3DL1+: control-2.1%, SLE-4.7%; KIR2DL3+: control-0%, SLE-0.9%; KIR2DL2+: control-0%, SLE-0.9%. (**D**) Expression of *KIR* transcripts (*KIR3DL1, KIR2DL3* and *KIR2DL2*) in synovial CD8^+^ T cells and expression of *FOXP3* in synovial CD4^+^ T cells from rheumatoid arthritis (RA, N=18) and osteoarthritis (OA, N=3) are shown. KIR3DL1+: OA-0.4%, RA-2.8%; KIR2DL3: OA-0%, RA-0.2%; KIR2DL2: OA-0%, RA-2.4%; FOXP3+: OA-9.6%, RA-9.4%.

Next, we investigated whether KIR^+^CD8^+^ T cells are also present in the inflamed tissues of patients with these diseases. We first analyzed the frequency of KIR^+^CD8^+^ T cells in the intestinal biopsies of CeD patients in remission or with active disease, as well as in healthy controls. Patients with active disease had higher levels of KIR^+^CD8^+^ T cells in the gut than those in remission (on a gluten-free diet) as well as the non-celiac controls (Fig. 1B), indicating a synchronous expansion of KIR^+^CD8^+^ T cells with incidence of autoimmune responses. In addition, we took advantage of publicly available single cell RNA-seq data from SLE kidneys and Rheumatoid Arthritis (RA) synovia previously generated by the Accelerating Medicines Partnership RA/SLE program (*13, 14*). A markedly increased number of CD8^+^ T cells expressing *KIR* transcripts (*KIR3DL1, KIR2DL3* and *KIR2DL2*) was observed in the kidneys of patients with SLE compared to healthy kidneys (Fig. 1C). In addition, we detected a higher frequency of KIR^+^CD8^+^ T cells in the synovial tissues of RA patients compared to those with Osteoarthritis (OA), while the percentages of synovial FOXP3^+^CD4^+^ regulatory T (Treg) cells were similar between RA and OA (Fig. 1D). Although both RA and OA cause joint inflammation, RA is a classic autoimmune disease (*15*), whereas OA is not, suggesting that induction of KIR^+^CD8^+^ T cells rather than CD4^+^ Tregs is more specific to autoimmune inflammations.

### KIR^+^CD8^+^ T cells are the functional and phenotypic equivalent of mouse Ly49^+^CD8^+^ T cells

Next we sought to investigate whether KIR^+^CD8^+^ T cells are the functional counterpart of mouse Ly49^+^ regulatory CD8^+^ T cells. Previously we found that Ly49^+^CD8^+^ T cells suppress myelin oligodendrocyte glycoprotein (MOG)-specific pathogenic CD4^+^ T cells in a perforin-dependent manner (*8*), as also suggested by Cantor and colleagues (*6*), indicating cytotoxicity as the mechanism of suppression. Deamidated gliadin derived from dietary gluten is the antigen for CD4^+^ T cells that drive autoimmune enteropathy in human CeD (*16, 17*). Therefore, we explored whether KIR^+^CD8^+^ T cells can suppress gliadin-specific CD4^+^ T cells from CeD patients. CD8^+^ T cells were purified from peripheral blood mononuclear cells (PBMCs) of HLA-DQ2.5^+^ CeD patients; from these KIR^+^CD8^+^ and KIR^-^CD8^+^ T cells were sorted, activated with anti-CD3/CD28 microbeads overnight, and then cultured with the CD8-depleted fraction of PBMCs at a 1:30 ratio in the presence of 250μg/mL deamidated gluten. The cultures were harvested on Day 6, and the gliadin-specific CD4^+^ T cells were enriched and quantified using PE-labeled HLA-DQ2.5 tetramers complexed with different gliadin peptides (Fig. 2A) (*18, 19*). In the absence of KIR^+^CD8^+^ T cells, deamidated gluten profoundly stimulated the expansion of gliadin-specific CD4^+^ T cells. Importantly, stimulated KIR^+^CD8^+^ T cells, but not KIR^-^CD8^+^ T cells or KIR^+^NK cells, significantly reduced the number of gliadin-specific CD4^+^ T cells (Fig. 2B and fig. S2B) without affecting the number of total CD4^+^ T cells (fig. S2A). This effect of KIR^+^CD8^+^ T cells appears to target only the pathogenic CD4^+^ T cells, since they had no discernible effect on hemagglutinin (HA)-specific CD4^+^ T cells induced by influenza A HA peptides (fig. S2C) or on proliferation of CD4^+^ T cells responding to anti-CD3 stimulation (fig. 2D). We also measured Annexin V binding on Day 3 (Fig. 2A) and found increased staining of gliadin-specific CD4^+^ T cells in the presence of KIR^+^CD8^+^ T cells (Fig. 2C), indicating these T cells induce apoptosis of the pathogenic CD4^+^ T cells. Since previous studies suggest that the regulatory function of mouse Ly49^+^CD8^+^ T cells is mediated by both recognition of H2-D^b^ (classical MHC I) (*8*) and Qa-1 (non-classical MHC I) (*5, 6, 9*) on their target cells, we investigated whether destruction of gliadin-specific CD4^+^ T cells by KIR^+^CD8^+^ T cells is dependent on their recognition of MHC I. Therefore, we blocked HLA-ABC or HLA-E with specific antibodies in the presence of KIR^+^CD8^+^ T cells and determined the frequency of gliadin-specific CD4^+^ T cells as a readout of the suppressive activity of KIR^+^CD8^+^ T cells. Our results show that anti-HLA-ABC blockade partially reversed the suppression by KIR^+^CD8^+^ T cells in 4/6 patients, while anti-HLA-E blockade restrained the suppressive functions of KIR^+^CD8^+^ T cells in 5/6 patients (p<0.05) (Fig. 2D). Taken together, our data suggest that KIR^+^CD8^+^ T cells specifically eliminate gliadin-specific pathogenic CD4^+^ T cells from the leukocytes of CeD patients via recognition of classical and/or non-classical MHC I molecules.

**Fig. 2.**
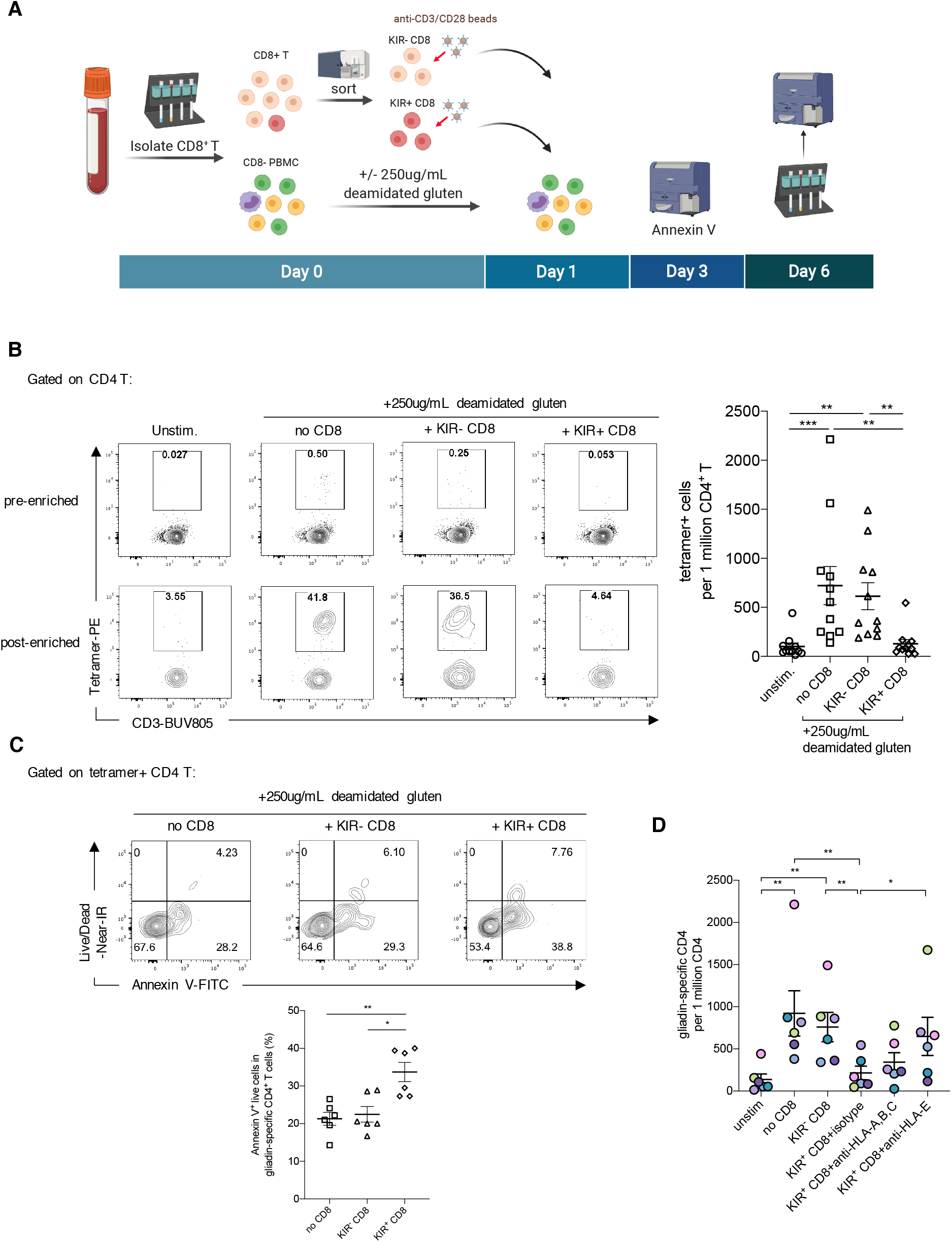
Elimination of gliadin-specific CD4^+^ T cells by KIR^+^CD8^+^ T cells. (**A**) Experimental schematic. (**B**) Representative contour plots showing the tetramer binding of CD4^+^ T cells after enrichment by MACS columns and summary of number of gliadin-specific CD4^+^ T cells (binding to HLA-DQ2.5 tetramers complexed with gliadin peptides) per 1 million CD4^+^ T cells on Day 6. Experiments were repeated using PBMCs from 11 CeD patients. \*\**P*<0.01, \*\*\**P*<0.001, Friedman test corrected for multiple comparisons. (**C**) Representative contour plots and summarized scatter plot displaying Annexin V binding of gliadins-specific (tetramer-positive) CD4^+^ T cells from the culture harvested on Day 3 (N=6). \**P*<0.05, \*\**P*<0.01, Friedman’s test corrected for multiple comparisons. (**D**) Summarized scatter plot displaying the frequency of gliadin-specific CD4^+^ T cells from the PBMC cultures in the presence or absence of pre-activated KIR^-^ or KIR^+^ CD8^+^ T cells with isotype control, anti-HLA-ABC or anti-HLA-E blockade. Results from 6 different CeD patients are indicated by different colors. \**P*<0.05, \*\**P*<0.01, Friedman test corrected for multiple comparisons.

To further investigate whether KIR^+^CD8^+^ T cells are the phenotypic equivalent of mouse Ly49^+^ T cells in humans, we performed RNA sequencing (RNA-seq) analysis on KIR^+^ versus KIR^-^ CD8^+^ T cells from patients with MS to compare with mouse Ly49^+^CD8^+^ T cells in EAE (mouse model of human MS). There were 778 differentially expressed genes (adjusted P<0.05, fold change >2) between KIR^+^ and KIR^-^CD8^+^ T cells, among them 300 were up-regulated and 478 were down-regulated in KIR^+^CD8^+^ T cells (table S1). Notably, KIR^+^CD8^+^ T cells showed a marked up-regulation of cytotoxic molecules (e.g. *GZMH, GZMB, PRF1* and *GNLY*), NK-associated genes (e.g. *NKG7, NCR1* and the *KLR* family) and cell-trafficking molecules (such as *CX3CR1*, which mediates migration of leukocyte to inflamed tissues and is involved in tissue injury-mediated brain inflammation, and brain-homing receptor *ITGB1*), in addition to inhibitory KIR receptor genes (Fig. 3A). In addition, KIR^+^CD8^+^ T cells had higher transcript levels for Helios (encoded by *IKZF2*), a transcription factor associated with regulatory functions of both CD4^+^ and CD8^+^ T cells (*7*). On the other hand, KIR^+^CD8^+^ T cells down-regulated naïve/memory T cell-associated molecules, e.g. *CCR7, SELL, TCF7* and *IL7R*, indicating they might have entered the program for effector T cell differentiation. Interestingly, KIR^+^CD8^+^ T cells had a lower expression of the co-stimulatory receptor *CD28* (Fig. 3A), which is one of the key features for regulatory CD8^+^ T cell populations in mice and humans (*20*). Gene Ontology enrichment analysis of these differentially expressed genes showed enrichment for T cell activation, proliferation, migration and differentiation (Fig. 3B). Moreover, gene set enrichment analysis (GSEA) (*21, 22*) revealed that about half of the top 200 genes up-regulated in Ly49^+^CD8^+^ T cells (including cytotoxic molecule *GZMB, KLRC* family genes, *CX3CR1, ITGB1* and *IKZF2*) were also higher in KIR^+^CD8^+^ T cells (Fig. 3C). Previously, we found Ly49^+^CD8^+^ T cells expressed 16 out of 60 of the genes conserved in CD4^+^ Tregs (*8*), and these same Treg signature genes (*23*) were also enriched in KIR^+^CD8^+^ T cells in GSEA analysis (Fig. 3D). Overall, RNA-seq analysis indicates that KIR^+^CD8^+^ T cells from MS patients share many similarities with Ly49^+^CD8^+^ T cells from EAE mice.

**Fig. 3.**
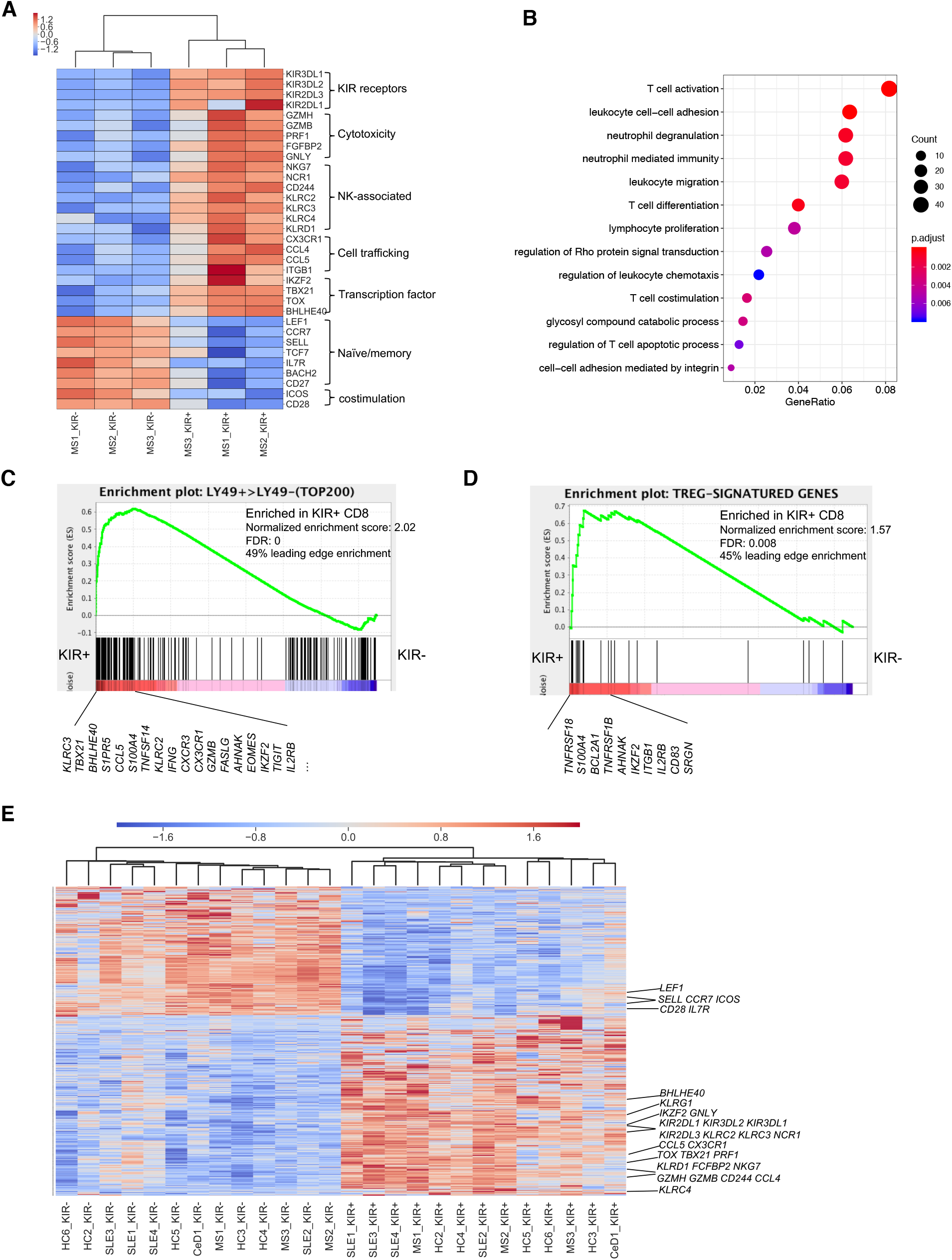
Bulk RNA-seq analysis of KIR^+^ versus KIR^-^ CD8^+^ T cells. (**A**) Heatmap displaying transcript levels of selected differentially expressed genes (DEGs) in KIR^+^ vs. KIR^-^ CD8^+^ T cells sorted from MS patients (N=3). (**B**) Gene Ontology (GO) enrichment analysis of DEGs between KIR^+^ and KIR^-^ CD8^+^ T cells from MS patients. **C**-**D**, Gene set enrichment analysis (GSEA) on KIR^+^ versus KIR^-^ CD8^+^ T cells from MS patients using the top 200 genes up-regulated in Ly49^+^ CD8^+^ T cells compared to Ly49^-^ cells (**C**) or signature genes of CD4^+^ regulatory T (Treg) cells (**D**) as reference gene sets. (**E**) Heatmap showing expression of the 963 DEGs in KIR^+^ and KIR^-^ CD8^+^ T cells from healthy subjects and patients with autoimmune diseases (MS, CeD or SLE) determined by bulk RNA-seq. Genes overlapping with DEGs defined in MS patients are annotated.

Furthermore, we also performed RNA-seq on KIR^+^ and KIR^-^CD8^+^ T cells from healthy subjects and patients with other autoimmune diseases, specifically CeD and SLE, to determine whether there are common features shared by KIR^+^CD8^+^ T cells across different circumstances. We identified a set of 963 genes that were differentially expressed (adjusted P<0.05, fold change >2) between KIR^+^ and KIR^-^CD8^+^ T cells from all subjects, including HC, MS, CeD, and SLE patients. Many of them overlapped with the differentially expressed genes previously defined in MS (Fig. 3E and table S2). However, larger fold changes of these genes were observed in patients with higher frequencies of KIR^+^CD8^+^ T cells (fig. S3A). Consistent with the transcriptional profiles, KIR^+^CD8^+^ T cells had higher protein expression levels for granzyme B, perforin, CX3CR1, KLRG1, CD244, TIGIT, T-bet and Helios proteins and lower levels of CCR7, CD27 and CD28, as measured by flow cytometry (fig. 3B). In addition, we compared KIR^+^ and KIR^-^ CD8^+^ T cells in kidneys or synovia for expression of the same genes enriched in circulating KIR^+^CD8^+^ T cells. Similar to those cells, both kidney and synovial KIR^+^CD8^+^ T cells up-regulated *KLRG1, CD244, TIGIT, CX3CR1, PRF1, GZMB* and *IKZF2*, while down-regulating *CD28* and *CCR7* (fig. S4). Overall, our results suggest KIR^+^CD8^+^ T cells are the functional and phenotypic equivalent of mouse Ly49^+^CD8^+^ T cells in humans, with conserved features in both healthy subjects and those with autoimmune diseases.

### Increased KIR^+^CD8^+^ T cells in SARS-CoV-2- and Influenza-infected patients

While previously it had been thought that most self-specific T cells were eliminated in the thymus, recent work shows that this is not the case, and that many such cells survive and populate the periphery of both humans and mice (*24, 25*). We have speculated that this is because the constant threat of infectious diseases throughout human history (*26*) necessitates a complete T cell repertoire (*24, 27*) such that even self-reactive T cells might be needed in the response to a particular pathogen. Consistent with this are classic experiments showing that infectious diseases or treatments that mimic them can activate self-specific T cells (*28, 29*). There is also anecdotal evidence that many patients with autoimmune diseases cite an infection immediately preceding the onset of their disease (*30-34*). Thus, we were interested in analyzing patients with an infectious disease to see whether KIR^+^CD8^+^ T cells were induced as part of the response. In particular, emerging reports show that infection with Severe Acute Respiratory Syndrome Coronavirus 2 (SARS-CoV-2) can lead to excessive production of pro-inflammatory cytokines and autoreactivity is one of the common features of severe disease with pathogenic prothrombotic autoantibodies detected in serum from patients hospitalized with Coronavirus Disease 2019 (COVID-19) (*35-39*). Therefore, we analyzed the frequency of KIR^+^CD8^+^ T cells in the peripheral blood of patients with COVID-19 as compared to age/gender-matched healthy subjects collected before the pandemic. The percentage of KIR^+^CD8^+^ T cells was elevated in COVID-19 patients and correlated with disease severity (Fig. 4A). Moreover, an even higher frequency of KIR^+^CD8^+^ T cells was found in COVID-19 patients with vasculitis or embolism and to a lesser extent in those with acute respiratory distress syndrome (ARDS) (Fig. 4B and figs. S5, C-D), which are common complications of this disease and likely caused by excessive inflammation. Therefore, increase of KIR^+^CD8^+^ T cells might be indicative for autoimmune-related immunopathology during SARS-CoV-2 infection. However, we did not observe a significant difference in the levels of CD25^hi^CD127^low^CD4^+^ Tregs or KIR^+^ NK in COVID-19 patients compared to healthy donors, or in COVID-19 patients with different disease severities or complications (figs. S5, A-D and Fig. 4B). We also utilized the publicly available single cell RNA-seq data of bronchoalveolar immune cells from COVID-19 patients and healthy people (*40*) to investigate whether KIR^+^CD8^+^ T cells are also present in the bronchoalveolar lavage fluid (BALF) of COVID-19 patients. Increased frequency of CD8^+^ T cells expressing *KIR* transcripts (*KIR3DL1, KIR3DL2, KIR2DL3* or *KIR2DL1*) were detected in the BALF of COVID-19 patients with moderate or severe disease compared to the BALF from healthy controls (Fig. 4C), which indicates KIR^+^CD8^+^ T cells are induced at the local site of infection as well. Additionally, in order to extend our findings in COVID-19 to other infectious diseases, we also measured the levels of KIR^+^CD8^+^ T cells in another infectious disease. In patients with acute influenza virus infection, we also observed an increased frequency of KIR^+^CD8^+^ T cells but not CD4^+^ Tregs in the peripheral blood of most patients (Fig. 4D), suggesting that KIR^+^CD8^+^ T cells are generally induced as part of the response during an infection.

**Fig. 4.**
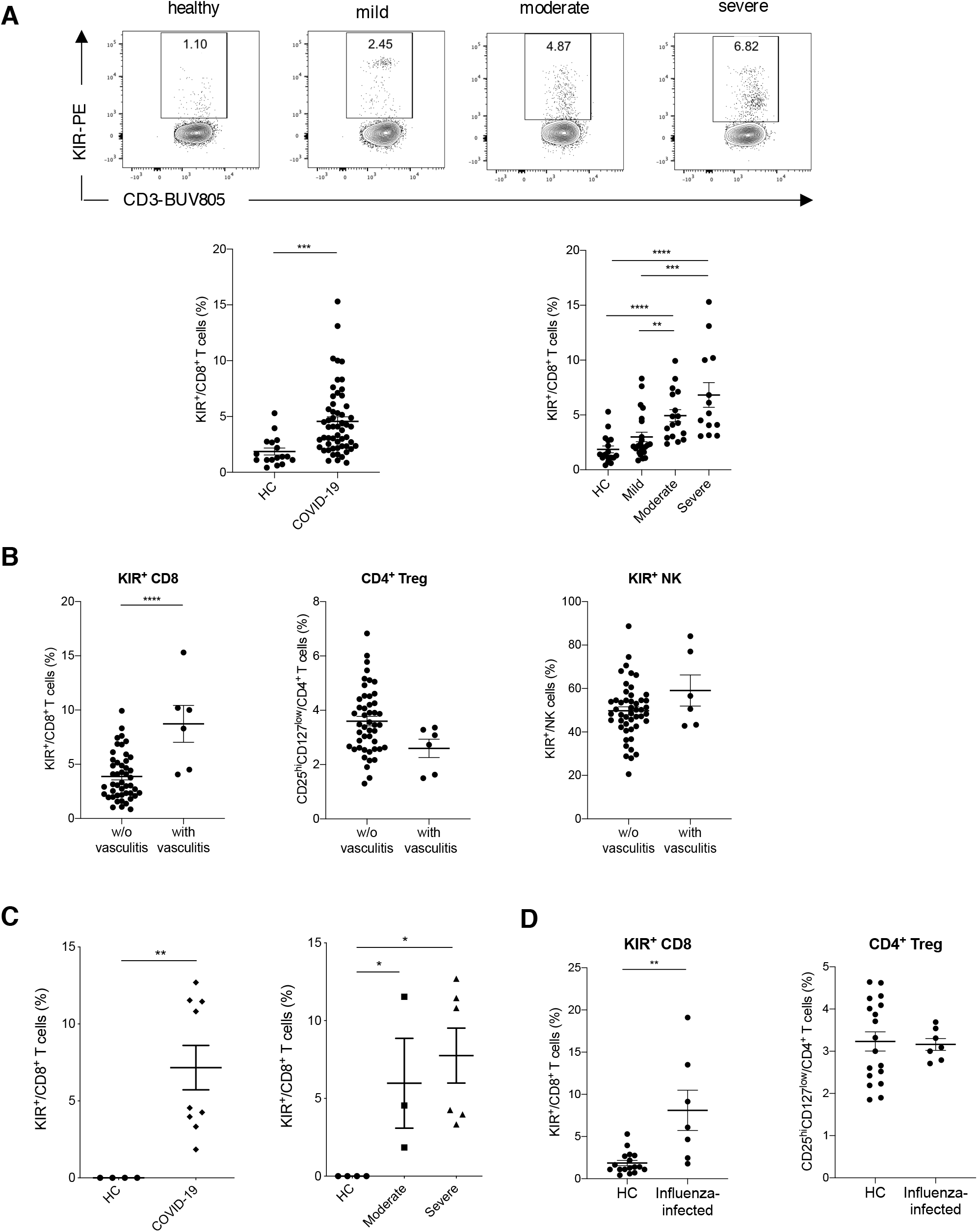
Increased KIR^+^CD8^+^ T cells in infectious diseases. (**A**) Representative contour plots and summarized scatter plots showing percentage of KIR^+^ cells in CD8^+^ T cells from the blood of 17 healthy controls and 53 COVID-19 patients with varying disease severity (mild: N=23, moderate: N=17, severe: N=13). left: \*\*\**P*<0.001, unpaired t test; right: \*\**P*<0.01, \*\*\**P*<0.001, \*\*\*\**P*<0.0001, Kruskal-Wallis test corrected for multiple comparisons. (**B**) Frequency of KIR^+^CD8^+^ T cells, CD4^+^ Tregs (CD25^hi^CD127^low^) and KIR^+^ NK cells in the blood of COVID-19 patients with or without vasculitis. \**P*<0.05, \*\*\*\**P*<0.0001, unpaired t test. (**C**) Frequency of CD8^+^ T cells expressing *KIR* transcripts (*KIR3DL1, KIR3DL2, KIR2DL3* or *KIR2DL1*) in the bronchoalveolar lavage fluid of healthy controls (N=4) versus COVID-19 patients (N=9) (left, \*\**P*<0.01, Mann-Whitney test) and healthy controls versus COVID-19 patients with moderate (N=3) or severe (N=6) disease (right, \**P*<0.05, Kruskal-Wallis test corrected for multiple comparisons). (**D**) Frequency of KIR^+^CD8^+^ T cells and CD4^+^ Tregs (CD25^hi^CD127^low^) in the blood of healthy controls (N=17) versus influenza-infected patients (N=7). \*\**P*<0.01, Mann-Whitney test.

### Commonality and heterogeneity of KIR^+^CD8^+^ T cells

In order to better understand the functional properties of this type of cells under different circumstances, we integrated the single cell RNA-seq data of peripheral blood CD8^+^ T cells from healthy subjects, MS patients and COVID-19 patients generated from the 10x Genomics platform (*41*) using the Seurat package (*42, 43*). Total CD8^+^ T cells were projected onto a two-dimensional UMAP and unsupervised clustering identified 8 subpopulations based on gene expression. KIR^+^CD8^+^ T cells from different conditions (healthy, MS and COVID-19) formed a distinct cluster with high expression of effector genes (*GZMB* and *PRF1*) as well as *KIR* transcripts (Figs. 5, A-B and table S3). When compared with KIR^-^ effector CD8^+^ T cells, KIR^+^ effector CD8^+^ T cells displayed higher expression of *IKZF2* (encoding the transcription factor Helios) and NK-associated genes (e.g., *TYROBP, KLRC2, KLRC3, NCR1* and *NCR3*), while showing an even lower expression of *IL7R, CD27* and *KLRB1*, in addition to marked expression of inhibitory KIR genes (table S3). These findings reveal the commonality of KIR^+^CD8^+^ T cells across physiological and diseased status as well as their uniqueness relative to other CD8^+^ T cells.

**Fig. 5.**
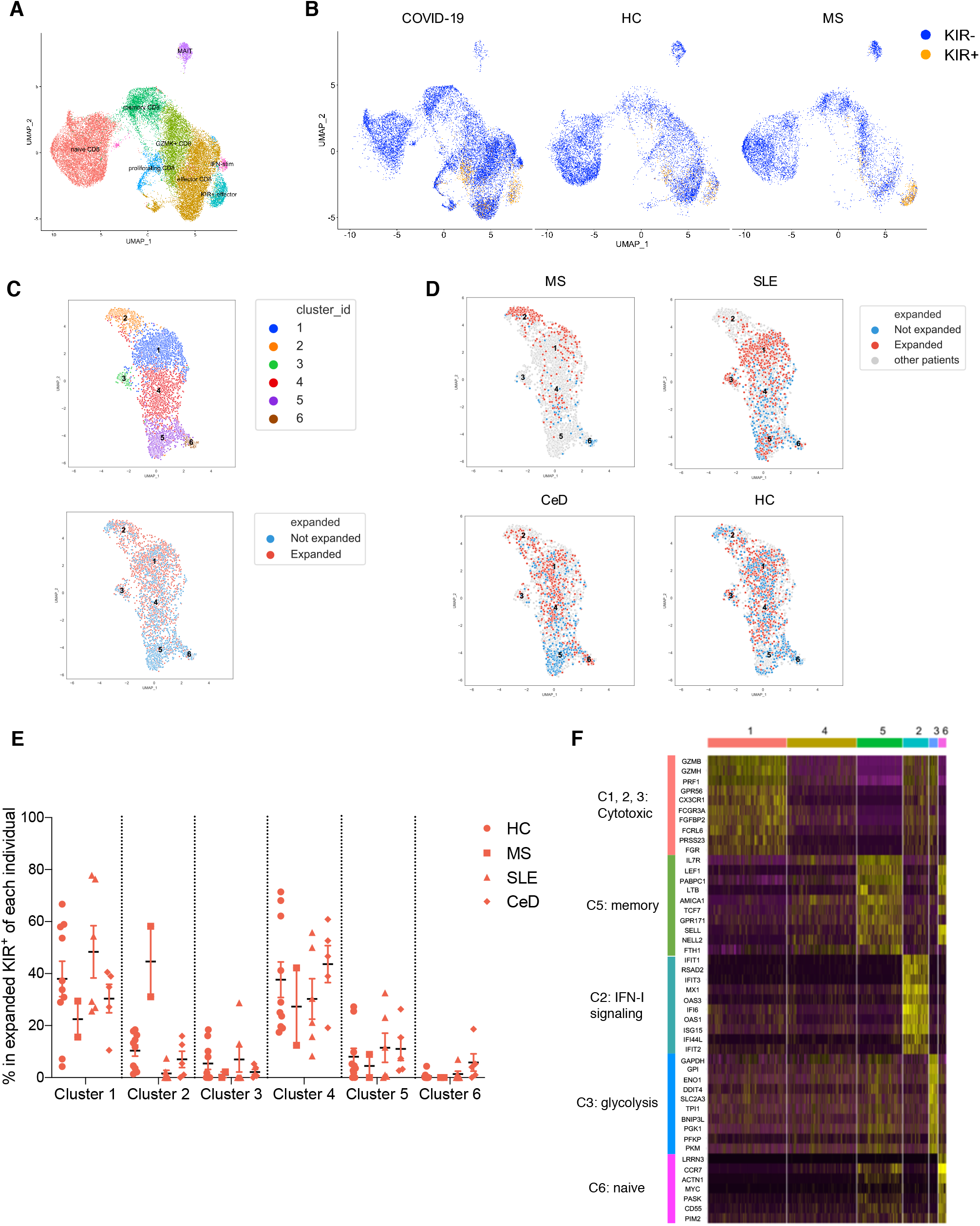
Single cell RNA-seq analysis of KIR^+^CD8^+^ T cells in the blood. **A**-**B**, Single cell RNA-seq analysis of total CD8^+^ T cells from the blood of healthy subjects (N=10), MS patients (N=6) and COVID-19 patients (N=25) by 10x Genomics. (**A**) UMAP plot of the 8 subpopulations identified by unsupervised clustering based on expression of marker genes in each cluster. (**B**) UMAP plots showing the distribution of KIR^+^CD8^+^ T cells (expressing *KIR3DL1, KIR3DL2, KIR2DL1* or *KIR2DL3* transcripts) and KIR^-^CD8^+^ T cells from healthy controls (HC), MS patients and COVID-19 patients. **C**-**F**, KIR^+^ CD8^+^ T cells in the blood of healthy controls (N=10) and patients with MS (N=2), SLE (N=6) and CeD (N=5) were sorted for single cell RNA-seq using the Smart-seq2 protocol and analyzed using the R package ‘Seurat’. (**C**) UMAP plots showing KIR^+^ CD8^+^ T cells segregated into 6 clusters (upper) and the distribution of expanded (≥2 cells expressing same TCR) and unexpanded (cell expressing unique TCR) cells (lower). (**D**) UMAP plots of KIR^+^ CD8^+^ T cells from MS, SLE, CeD and HC are shown, with expanded and unexpanded cells annotated with different colors (expanded: red, unexpanded: blue, other diseases: grey). (**E**) Cluster compositions of expanded KIR^+^ CD8^+^ T cells from each individual. (**F**) Heatmap showing expression of the top 10 genes differentially expressed in each cluster, with the categories of each group of genes annotated on the left.

In order to better understand the similarity and heterogeneity of KIR^+^CD8^+^ T cells under different circumstances and to probe the mechanism for their suppressive activity to pathogenic CD4^+^ T cells, we performed single cell RNA-seq on 4,512 KIR^+^CD8^+^ T cells sorted from the blood of age- and gender-matched healthy subjects (N=10) and patients with MS (N=2), SLE (N=6) or CeD (N=5) using the Smart-seq2 protocol (*44*). In parallel, we also analyzed their T-cell receptor (TCR) α and β sequences (*45*). Unsupervised clustering of these KIR^+^CD8^+^ T cells by Seurat identified 6 clusters, with Clusters 1 to 3 mostly containing expanded KIR^+^CD8^+^ T cells (≥2 cells expressing same TCR) and Clusters 5 and 6 consisting of unexpanded cells expressing unique TCRs (Fig. 5C and figs. 6, A-B). Expanded KIR^+^ cells in Clusters 1 to 3 had higher transcripts for cytotoxic molecules (e.g., *GZMH, GZMB* and *PRF1*) and genes associated with effector T cells (e.g., *FCGR3A, FGFBP2* and *CX3CR1*). Cluster 2, which was more restricted to expanded KIR^+^ cells from MS patients, showed higher levels of Type I IFN responding genes (including *IFIT1, IFIT2, IFIT3, MX1, RSAD2* and *ISG15*). Cluster 3, specific to expanded KIR^+^ cells from a subset of HC and SLE patients, displayed higher expression of genes involved in glycolysis (e.g., *GAPDH, GPI, ENO1* and *PGK1*) (Figs. 5, D-F and table S4). Cells in Cluster 4 were in a transitional state with a loss of memory-associated features. Clusters 5 and 6 (restricted to unexpanded KIR^+^CD8^+^ T cells) displayed memory and naïve signatures, respectively (Fig. 5F and table S4), and accounted for a small proportion of total KIR^+^CD8^+^ T cells (fig. S6A). T cell clones expressing identical TCRs can be found in different clusters, indicating possible lineage relationships. In addition, clonally expanded KIR^+^CD8^+^ T cells in COVID-19 patients identified from the previous 10x Genomics scRNA-seq (*46*) displayed a higher expression of cytotoxic genes while down-regulating naïve- or memory-associated genes compared to unexpanded KIR^+^CD8^+^ T cells (figs. S6, C-D). Thus, in parallel with clonal expansion, KIR^+^CD8^+^ T cells might lose their naïve or memory attributes, enter the differentiation program for effector T cells and then suppress pathogenic CD4^+^ T cells via cytotoxicity. There are common features shared by KIR^+^CD8^+^ T cells from healthy subjects and different diseases, yet there is also heterogeneity (i.e., up-regulated Type I IFN signaling and glycolysis in Clusters 2 and 3) associated with different diseases or treatments.

**Fig. 6.**
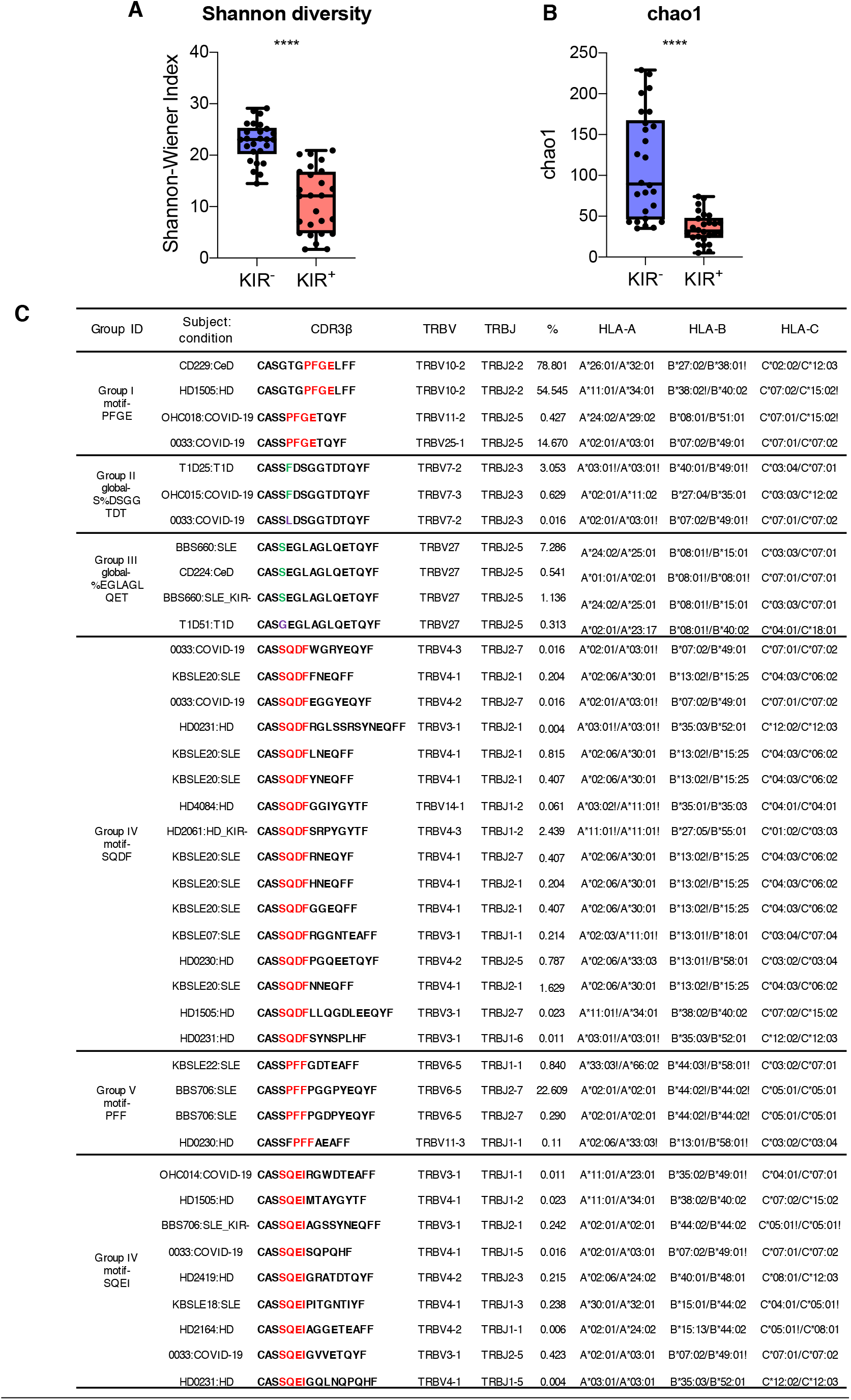
Analysis of T cell receptor sequences of KIR^+^CD8^+^ T cells. **A**-**B**, Summary histograms showing Shannon-Wiener Index (**A**) and chao estimates (**B**) of TCRs of KIR^-^ versus KIR^+^ CD8^+^ T cells from 26 subjects including 11 healthy donors, 2 MS, 5 SLE, 3 CeD and 5 T1D patients as evaluated by VDJtools. \*\*\*\**P*<0.0001, Wilcoxon matched-pairs signed rank test. (**C**) Selected specificity groups from the GLIPH2 analysis that contained TCRs from 3 or more individuals and exhibited significant bias of V-gene usage (P<0.05) are shown.

To get a comprehensive understanding on the TCR repertoire of KIR^+^CD8^+^ T cells, we used the VDJtools (*47*) to estimate the diversity of KIR^+^CD8^+^ TCRs as compared to KIR^-^CD8^+^ TCRs from the same individuals (N=26). We found TCRs of KIR^+^CD8^+^ T cells had significantly lower Shannon-Wiener Index and chao estimates than that of KIR^-^CD8^+^ T cells (Figs. 6, A-B), indicating the TCR repertoire of KIR^+^CD8^+^ T cells is less diverse, which is also consistent with the previous report that KIR^+^CD8^+^ T cells display a more restricted TCR Vβ chain usage (*11*). In order to compare the antigen specificities of KIR^+^CD8^+^ T cells from different disease types, we also utilized GLIPH2 (*48*), which is an algorithm to cluster TCRs that recognize the same antigen in most cases, to analyze single-cell TCR sequences of sorted KIR^+^ (6815 unique TCRs) or KIR^-^ CD8^+^ T cells (1630 unique TCRs) from healthy controls (N=10), MS (N=2), SLE (N=20), CeD (N=5), T1D (N=5) and COVID-19 (N=5) patients and bulk TCRβ sequences of sorted KIR^+^CD8^+^ T cells from 9 healthy controls (5607 unique TCRβ) along with their class I HLA alleles. The GLIPH2 analysis generated 982 clusters and 668 of them were shared between any two sources. We further filtered the resulting GLIPH clusters to 62 specificity groups that contained TCRs from 3 or more individuals and exhibited significant bias of V-gene usage (P<0.05) and some of them are shown in Figure 6C. TCRs of KIR^+^CD8^+^ T cells from healthy donors and patients with autoimmune diseases or COVID-19 can be grouped into the same GLIPH clusters, although with different extent of clonal expansion, indicating they might recognize same antigens commonly exist under physiological and different pathological conditions. Therefore, expanded KIR^+^CD8^+^ T cells have shared phenotypes and antigen specificity independent of the disease types studied here. While the analysis above shows that the TCR repertoire of KIR^+^CD8^+^ T cells is less diverse than CD8^+^ T cells generally, it is still considerably diverse, utilizing multiple classical HLAs and HLA-E, and probably many antigenic peptides as well. Previously we have found an average of 5 GLIPH clusters for each peptide MHC ligand (*49*), so there may be 10 or more such ligands for these T cells at a minimum. The identification of these ligands, and whether they extend beyond pathogenic CD4^+^ T cells will be important to determine in the future.

## Discussion

Here we characterize KIR^+^CD8^+^ T cells as a novel regulatory CD8^+^ T cell subset in humans, which suppress pathogenic CD4^+^ T cells in CeD, and likely other autoimmune diseases or infectious diseases as well, via their cytolytic activity. Similar to the perforin- or Fas/FasL-dependent suppression of self-reactive CD4^+^ T cells by murine Ly49^+^CD8^+^ T cells (*8, 50*), human KIR^+^CD8^+^ T cells might also target pathogenic CD4^+^ T cells via their cytolytic activity, since expanded KIR^+^CD8^+^ T cells significantly up-regulated cytotoxic molecules and increased apoptosis in gliadin-specific CD4^+^ T cells was observed in the presence of KIR^+^CD8^+^ T cells. This effect of KIR^+^CD8^+^ T cells is specific to self-reactive or otherwise pathogenic T cells, but not CD4^+^ T cells recognizing foreign antigens, since presence of KIR^+^CD8^+^ T cells had no discernable effects on HA-specific CD4^+^ T cells. In addition, the destruction of pathogenic CD4^+^ T cells by KIR^+^CD8^+^T cells appears to depend on recognition of both classical and non-classical I MHC molecules, since the blockade of either HLA-ABC or HLA-E can reverse the suppression by KIR^+^CD8^+^ T cells. We often observe an increased frequency of KIR^+^CD8^+^ T cells in the blood and also in the inflamed tissues of patients with autoimmune disease, and this positively correlates with disease activity of CeD. This expansion of KIR^+^CD8^+^ T cells correlating with the incidence of autoimmune responses might act as a negative feedback mechanism to ameliorate pathogenesis by killing autoreactive T cells. Besides, increased KIR^+^CD8^+^ T cells were found in COVID-19 patients and were associated with autoimmune-related complications. These increases were not restricted to SARS-CoV-2 infection, but also observed in patients with influenza virus infection, indicating increase of KIR^+^CD8^+^ T cell is a general mechanism induced during an infection. Therefore, we hypothesize that a primary role of KIR^+^CD8^+^ T cells might be to control pathogenic T cells that arise from self-reactivity in autoimmune disorders or in the course of an infectious disease owing to their cross-reactivity to antigens expressed by a particular pathogen. This would allow an organism to maintain as complete a peripheral T cell repertoire as possible to protect itself against potential infection by pathogens, yet still be able to precisely control T cell clones with cross-reactivity to self antigens. Interestingly, clonally expanded KIR^+^CD8^+^ T cells are also found in the peripheral blood of healthy subjects and share gene expression signatures as well as some antigen specificities with those from patients with autoimmune diseases or COVID-19. This indicates that at least some T cells of this type are continually active-although not at the very high levels that we see in COVID-19 patients or in some subjects with autoimmunity. This suggests that the activation of KIR^+^CD8^+^ T cells is a specific regulatory mechanism to maintain peripheral tolerance, even in healthy people. This type of peripheral tolerance is distinct from and likely complementary to CD4^+^ Tregs, which represent a separate lineage of T cells and does not appear to be generally active in the human infectious diseases analyzed here or murine infection (*51, 52*). Thus, the KIR^+^CD8^+^ T cells represent an important new factor in understanding peripheral tolerance and the relationship between autoimmunity and infectious diseases. Further characterization of this pathway and how it may be breaking down in autoimmune diseases and severe infections like COVID-19, will be important challenges for the future work. Our findings on the KIR^+^CD8^+^ T cells and their properties described here are likely to be useful in understanding key cellular dynamics in immune dysregulation and in potential therapeutic approaches to suppress undesirable self-reactivity in autoimmune or infectious diseases.

## Acknowledgments

We thank members of Davis lab for helpful discussions. We also thank J. Coller in Stanford Functional Genomics Facility for NGS sequencing (NIH award S10OD018220); the Stanford Protein and Nucleic Acid Facility for performing fragment analysis; Rahul Sinha, S. D. Conley and I. L. Weissman for assistance in single cell RNA-seq library preparation; the Stanford Shared FACS facility for assistance in flow cytometric analysis and cell sorting; L. Chen for help in python scripts; D. Chen in Institute for Systems Biology, Seattle for help with the COVID-19 10x single-cell RNA-seq data processing; R. Wittman and R. Puri from the Stanford occupational health clinic and I. Chang, J. Krempski, E. Do, M. Manohar, A. Fernandez, W. Zhang for collecting and processing blood samples from COVID-19 patients; Z. He and J. Fitzpatrick for data management and entry of COVID-19 patients; D. Furman, B. Gaudillière and D. Feyaerts for their help in obtaining COVID-19 PBMCs.

## Author contributions

J.L., N.S. and M.M.D conceived the study. J.L. and N.S. performed the experiments. M.Z., J.L. S.H.C., J.P. and A.T.S. analyzed the single cell RNA-seq data. M.J., L.C. and L.M.Steinmetz contributed to analysis of bulk RNA-seq data. X.J. prepared the bulk RNA-seq libraries. A.C. and L.M.Sollid provided the HLA-DQ2.5 molecules. Jiefu Li prepared samples for bulk TCRβ sequencing. J.W. and A.M.M. performed single cell TCR-seq. Y.S. and J.R.H. generated the single cell RNA-seq data of COVID-19 patients. V.V.M. prepared the HLA-DR4 molecules. B.A.P. and C.K. provided deamidated gluten. L.B.K., J.E.D., S.L.H. and J.R.O. provided PBMCs from MS patients. K.B., W.H.R. and P.J.U. provided PBMCs from SLE patients. K.C.N. and N.Q.F. provided blood and biopsy samples from CeD patients. K.C.N. and G.K.R.D. provided PBMCs and clinical metadata of COVID-19 patients. J.L., N.S. and M.M.D. wrote the manuscript with inputs from all the authors.

## Data and materials availability

Data generated in this study are included within the paper or are available from the corresponding authors upon reasonable request. Raw RNA-seq data that support the findings of this study have been deposited in the Gene Expression Omnibus under the accession number GSE168527. The code used for data analyses in this study are available from the corresponding authors upon reasonable request.

## Supplementary Materials

Materials and Methods

Supplementary Text

Figs. S1 to S6

References (*53*–*64*)

Tables S1 to S6

