## Supplementary Materials for "Human KIR^+^CD8^+^ T cells target pathogenic T cells in Celiac disease and are active in autoimmune diseases and COVID-19"

### Materials and Methods

#### Human samples

Our study cohort of patients with autoimmune disorders met classification criteria for systemic lupus erythematosus (SLE) (*53*), celiac diseases (CeD) (*54*) or Multiple sclerosis (MS) (*55*), respectively. Collection of blood or biopsies from patients with SLE, CeD or MS was covered by IRB-14734 (Stanford University Immunological and Rheumatic Disease Database: Disease Activity and Biomarker Study), IRB-20362 (Studying the Molecular Factors Involved in Celiac Disease Pathogenesis) and IRB-8629 (Understanding Mechanisms of Allergy and Immunology Study). Blood samples from patients during influenza virus infection were obtained from patients who had influenza-like symptoms and were tested positive for influenza A virus at the Emergency Department or the Express Outpatient Clinic at Stanford Hospital, which is covered by IRB-22442 (Immune Responses to Influenza-like Illness). Blood from healthy subjects was requested from Stanford Blood Center or drawn from healthy volunteers under IRB-40146. The protocols mentioned above have been approved by the Research Compliance Office of Stanford University. PBMCs from MS patients were also obtained from the Multiple Sclerosis Center at the University of California, San Francisco (UCSF) with the protocol approved by the committee on Human Research at UCSF. Informed written consent was obtained from all participants. Detailed information of the healthy controls and patients with autoimmune diseases included in the study is provided as Table S5. PBMCs were isolated from the blood through density gradient centrifugation (Ficoll-Paque, GE Healthcare). Duodenal biopsies from CeD patients were treated twice with 6 mM EDTA in calcium/magnesium-free HBSS for 30 min at 37 °C. Supernatants containing the epithelial fractions were combined, washed and kept on ice until staining. The remaining tissues were minced and incubated with 200 µg/mL Liberase TL and 20 U/mL DNase

I in IMDM for 30 min at 37 °C. Digested cell suspension was passed through 100 μm cell strainer, washed with complete media and combined with the epithelial fraction for staining.

COVID-19 patients and sample collection: enrollment included any adult with RT-PCR positive COVID-19. Informed consent was obtained from each patient or from the patient's legally authorized representative if the patient was unable to provide consent. Participants were excluded if they were taking any experimental medications (i.e. those medications not approved by a regulatory agency for use in COVID-19). COVID-19 severity of illness was defined as described in the literature (*56*). Collection of blood from COVID-19 patients was covered by IRB-14734 and NCT04373148. Handling of COVID-19 PBMCs for flow cytometric analysis was covered under APB-3343-MD0620. The IRB and APB protocols mentioned above have been approved by the Research Compliance Office of Stanford University. Clinical metadata was obtained from Stanford clinical data electronic medical record system as per consented participant permission and definitions and diagnoses of disease were used according to Harrison's Principles of Internal Medicine, 20e. Clinical metadata for the COVID-19 patients in this study is presented in Table S6.

**Flow cytometric analysis**

The following fluorescent dye-conjugated anti-human antibodies were used for staining: CD8a (RPA-T8), CD56 (5.1H11), TCR γ/δ (B1), KIR3DL1 (Dx9), KIR2DL2/L3 (Dx27), KIR2DL5 (UP-R1), TIGIT (A15153G), KLRG1 (SA231A2), CD244 (C1.7), CX3CR1 (2A9-1), CD28 (CD28.2), CD27 (O323), CCR7 (G043H7), T-bet (4B10), Helios (22F6), Granzyme B (QA16A02), Perforin (B-D48) and CD4 (RPA-T4) (Biolegend); CD3 (UCHT-1) and TCRαβ (IP26) (BD); KIR2DL1 (clone#143211) and KIR3DL2 (clone#539304) (R&D). Frozen cell samples were thawed and washed in 10% FBS with Benzonase (Sigma-Aldrich, 1:10,000) in RPMI. After 450g centrifugation, cells were treated with 1:20 diluted FcR block (Biolegend) in

FACS buffer (0.5% BSA, 2 mM EDTA in PBS) for 10 min followed by staining with antibodies against surface molecules (30 min, 4 °C). For intracellular staining, cells were fixed and permeabilized with the Intracellular Fixation & Permeabilization Buffer Set (eBioscience), followed by staining with antibodies against intracellular antigens (30 min, 4 °C). Cells were acquired on an LSR II flow cytometer (BD), and data was analyzed using FlowJo X. Dead cells were excluded based on viability dye staining (LIVE/DEAD™ Fixable Near-IR Dead Cell Stain, ThermoFisher).

**Functional assay**

Chymotryptic gluten digests were deamidated with recombinant human transglutaminase 2, as described previously (*57*). PBMCs were isolated from blood of HLA-DQ2.5$^{+}$ CeD patients on Day 0. CD8$^{+}$ T cells were purified from PBMCs using CD8 microbeads (Miltenyi) per manufacturer's instructions, stained with flow antibodies, and live CD3$^{+}$CD56$^{-}$CD8$^{+}$KIR$^{+}$ or KIR$^{-}$ T cells were sorted out by FACSAria Fusion flow cytometer (BD). The sorted KIR$^{+}$ and KIR$^{-}$ CD8$^{+}$ T cells were stimulated with anti-CD3/CD28 beads (Gibco) at 1:1 ratio (1 µL beads per $4\times10^{4}$ cells) supplemented with 50 U/mL IL-2 in 96-well plates for 18 hours. KIR$^{+}$ and KIR$^{-}$ NK cells were sorted for PBMCs and rested overnight. The CD8$^{-}$ PBMCs were stimulated with 250 µg/mL deamidated gluten or 10ug/mL Influenza A HA 306-318 peptide (PKYVKQNTLKLAT) or left unstimulated at $3\times10^{5}\sim1\times10^{6}$/100 µL per well supplemented with 50 U/mL IL-2. X-VIVO 15 with Gentamicin L-Gln (Lonza) supplemented with 10% human AB serum (Sigma-Aldrich) was used as culture medium. After 18 hours, anti-CD3/CD28 beads were removed from CD8$^{+}$ T cells by a magnet and KIR$^{+}$ or KIR$^{-}$ CD8$^{+}$ T cells or NK cells were added to the culture of CD8$^{-}$ PBMCs at 1:30 ratio. In the MHC I blockade experiments, 10 µg/mL anti-HLA-ABC (W6/32, Biolegend), anti-HLA-E (3D12, eBioscience) or isotype controls were added to the culture. 50 U/mL IL-2 was added to the cultures on Day 3. Cells were harvested on Day 6 and stained with 10 μg/mL HLA-DQ2.5 tetramers (*19*) complexed with four disease-relevant and immunodominant gliadin T cell epitopes (DQ2.5-glia-α1a, QLQPFPQPELPY; DQ2.5-glia-α2, PQPELPYPQPE; DQ2.5-glia-ω1, QQPFPQPEQPFP; DQ2.5-glia-ω2, FPQPEQPFPWQP) (*18*) or 10 μg/mL HLA-DR4 tetramers complexed with Influenza A HA 306-318 peptide for 45 min at room temperature. Magnetic bead enrichment of tetramer-positive $CD4^+$ T cells was done as previously described (*58*). Cells were washed with FACS buffer and then stained with antibodies against surface molecules for 30 min at 4 °C. After two washes with FACS buffer, 10% of the cells were reserved for FACS analysis while 90% were labeled with anti-PE microbeads and subjected to magnetic bead enrichment of PE-conjugated tetramer-positive cells using a single MACS column according to the manufacturer's protocol (Miltenyi). Cells were also harvested on Day 3 to measure Annexin V binding (BD) on gliadin-specific $CD4^+$ T cells. All cells were acquired on an LSR II flow cytometer (BD), gated on live $CD3^+CD4^+CD8^-TCR\alpha\beta^+$ cells, and analyzed using FlowJo X software. The frequency of tetramer-positive cells was calculated by dividing the number of post-enrichment $tetramer^+$ $CD4^+$ T cells by the number of $CD4^+$ T cells in the pre-enrichment sample multiplied by 9 (to account for the fact that 90% of the cells were used for the enrichment).

**Bulk RNA-seq gene expression quantification and data analysis**

Bulk RNA sequencing was done as previously described (*59*). Live $KIR^+$ or $KIR^-$ $CD8^+$ T cells were bulk sorted directly into 350 μL TRIzol (Qiagen) by FACSAria Fusion flow cytometer (BD). Total RNA was extracted from TRIzol samples using chloroform separation and isopropanol precipitation and then RNAeasy Plus Mini kit (Qiagen) for clean-up. After analysis on the 2100 Bioanalyzer, the sequencing libraries were prepared using the RiboGone Mammalian rRNA Depletion Kit (Clontech) and the SMARTer Stranded RNA-seq Kit (Clontech). The resulting library was sequenced on the HiSeq 4000 platform (Illumina) in Stanford Functional Genomics Facility. For each sample in the bulk RNA sequencing library, 75-base-pair paired-end reads were acquired from the sequencer. We aligned the reads to the human reference genome (NCBI GRCh38) using STAR v2.7.0e (*60*). Gene counts were quantified and normalized (TPM) with Salmon (*61*). Differential gene expression analysis was determined via the DESeq function in the DESeq2 R package (*62*). Heatmaps were generated with seaborn.clustermap in python. Gene Ontology analysis plots were generated with the R package 'clusterProfiler'. To generate gene sets for gene set enrichment analysis (GSEA), we selected the top 200 genes up-regulated in $Ly49^+$ $CD8^+$ T cells compared to $Ly49^-$ $CD8^+$ T cells in EAE mice (*8*), and the previously reported $CD4^+$ Treg signature genes identified in mice (*23*). These mouse genes were converted to homologue genes in humans and constituted as gene sets for the subsequent GSEA analysis (*21, 22*) in human $KIR^+$ versus $KIR^-$ $CD8^+$ T cells.

**Analysis of single cell RNA-seq of kidneys and synovial tissues**

The Unique Molecular Identifier (UMI) count matrixes of cells in kidneys (accession code SDY997) (*13*) or synovial tissues (accession code SDY998) (*14*) generated by CEL-Seq2 were downloaded from the ImmPort repository and downstream analysis was performed using the Seurat 3.0 package. Cells with fewer than 1,000 detected genes, more than 5,000 detected genes or more than 25% mitochondrial genes were discarded. $CD8^+$ T cells (expressing *CD3D*, *CD3E*, *CD8A* and *CD8B* transcripts) and $CD4^+$ T cells (expressing *CD3D*, *CD3E* and *CD4* transcripts) were selected for standard downstream procedures of log-normalization, variable gene selection and data scaling.

**Analysis of single cell RNA-seq of bronchoalveolar immune cells**

Filtered expression matrix of single-cell RNA-seq of immune cells from the bronchoalveolar lavage fluid of 6 severe and 3 moderate COVID-19 patients and 3 healthy controls generated by 10x Genomics (*40*) were downloaded from Gene Expression Omnibus under the accession number GSE145926. $CD8^+$ T cells were identified for downstream analysis using the Seurat 3.0 package.

**Analysis of single cell RNA-seq generated by 10x Genomics**

Single cell RNA-seq of T cells from the blood of healthy subjects (N=10), MS patients (N=6) and COVID-19 patients (N=25)(*46*) from the microfluidic droplet platform (10x Genomics Chromium Single Cell 5' paired-end chemistry) were de-multiplexed, aligned to the GRCh38 reference genome, and converted into gene counts matrices using CellRanger 3.1.0. Downstream analysis was performed using the Seurat 3.0 package. Cells with fewer than 800 detected genes, more than 3,000 detected genes or more than 10% mitochondrial genes were discarded. $CD8^+$ T cells (expressing *CD8A* and *CD8B* but not *TRDC* transcripts) from each individual were selected for further analysis. To make counts comparable among cells, gene counts were normalized to 10,000 reads per cell, then log-transformed. We identified highly-variable genes for each individual, then integrated gene expression data from all individuals using Seurat's integration anchor discovery algorithm (*43*). We performed PCA dimensionality reduction on the integrated data, then clustered cells with the Louvain algorithm and visualized the data using UMAP. We identified canonical cell type marker genes that were conserved across conditions using the Wilcoxon rank-sum test implemented in the Seurat package's 'FindConservedMarkers' function.

**Single cell RNA-seq gene expression quantification by Smart-seq2 and data analysis**

Single cell RNA-seq of blood $KIR^+$ $CD8^+$ T cells (Live $CD3^+CD56^-CD8^+TCR\alpha\beta^+KIR^+$ cells) was performed using the Smart-seq2 protocol with some modifications (*44, 63*). In brief, single cells were sorted into 96-well plates containing 5 μL lysis buffer (0.8 U/μL RNase Inhibitor (Clontech),

~5,000 molecules of ERCC (External RNA Controls Consortium) spike-in RNAs (Ambion), 0.08% BioUltra Triton X-100 (Sigma-Aldrich), 2 μM oligo-d$T_{30}$VN (Integrated DNA Technologies, 5′-AAGCAGTGGTATCAACGCAGAGTAC$T_{30}$VN-3′), 2 mM Qiagen dNTP mix) in each well. Immediately after sorting, plates were sealed with aluminium seal (Axygen), centrifuged, flash frozen on dry ice and then stored at -80 °C. Before reverse transcription, the plates were thawed on ice and lysed at 72 °C for 3 min. 5 μL reaction mix containing 10 mM DTT, 2 μM TSO (Exiqon, 5'-AAGCAGTGGTATCAACGCAGAGTGAATrGrGrG-3′), 20 U/μL SMARTScribe Reverse Transcriptase (Takara), 2 U/μL RNase Inhibitor (Clontech) and 2× First Strand Buffer was added to each well and reverse transcription was carried out by incubating wells on a thermal-cycler (Eppendorf) at 42 °C for 90 min, 10 cycles of 50 °C for 2 min, 42 °C for 2 min, and stopped by heating at 70 °C for 15 min. Subsequently, 15 μL of PCR mix containing 1.67× KAPA HiFi HotStart ReadyMix (Roche, KK2602) and 0.17 μM IS PCR primer (IDT, 5′-AAGCAGTGGTATCAACGCAGAGT-3′) was added to each well and second-strand synthesis was performed on a thermal-cycler (Eppendorf) by using the following program: 1) 98 °C for 3 min, 2) 22 cycles of 98 °C for 20 s, 67 °C for 15 s and 72 °C for 6 min, and 3) 72 °C for 5 min. 1 μL of the cDNA products were used for TCR PCR reaction. The remaining 24 μL cDNA products were subjected to purification by AMPure XP beads (Beckman Coulter) on the Biomek FX$^P$ Automated Workstation (Beckman Coulter): 15.6 μL of Ampure XP beads (0.65×) were added to each sample and mixed by pipetting up and down thirty times; the mixture were incubated at room temperature for 5 min to let the DNA bind to the beads; then the 96-well plate was placed on the magnet for 5 min, and the liquid was removed while samples were on the magnet; the beads were wash with 180 μL of 80% (vol/vol) ethanol solution twice and air dried on the magnet for 6 min; 25 μL of water was added to each well, mixed by pipetting up and down ten times, and incubated at room temperature for 3 min; the plate was placed on the magnet for 3 min and the supernatants were transferred to a new 96-well plate; finally, 2 μL of the supernatants were subjected to quality control using capillary electrophoresis on a Fragment Analyzer (Agilent Technologies) by Stanford Protein and Nucleic Acid Facility.

cDNA in 96-well plates was transferred into 384-well Low Volume Serial Dil. (LVSD) plates (TTP Labtech) and diluted to 0.16 ng/μL using a Mosquito X1 liquid handler (TTP Labtech). Illumina sequencing libraries were prepared as described previously(*64*) using a Mosquito HTS liquid handler (TTP Labtech). In brief, tagmentation was carried out on 0.4 μL double-stranded cDNA using the Nextera XT DNA Library Preparation Kit (Illumina, FC-131-1096). Each well was mixed with 0.8 μL Nextera tagmentation DNA buffer (Illumina) and 0.4 μL Amplicon Tagment Mix (Illumina), then incubated at 55 °C for 10 min. The reaction was stopped by adding 0.4 μL Neutralize Tagment Buffer (Illumina) and centrifuging at room temperature at 3,000 g for 5 min. Indexing PCR reactions were performed by adding 0.8 μL of pre-mixed 5 μM i5 and i7 unique dual indexing primers (IDT, customized) and 1.2 μL of Nextera NPM mix (Illumina). PCR amplification was carried out on a C1000 Touch™ Thermal Cycler with 384-Well Reaction Module (Bio-rad) using the following program: 1) 72 °C for 3 min, 2) 95 °C for 30 s, 3) 12 cycles of 95 °C for 10 s, 55 °C for 30 s and 72 °C for 1 min, and 4) 72 °C for 5 min.

After library preparation, wells of each library plate were pooled using a Mosquito HTS liquid handler (TTP labtech). Pooling was followed by two purifications using 0.65× and 1× AMPure XP beads (Beckman Coulter), respectively. Library quality was assessed by Agilent 2100 Bioanalyzer and normalized to 5 nM. Libraries were sequenced on the Hiseq4000 Sequencing System (Illumina) in Stanford Functional Genomics Facility, acquiring 150-bp paired-end reads.

Stanford Functional Genomics Facility extracted and generated FASTQ files for each cell, distinguished by the unique dual index adaptors. Reads were aligned to the GRCh38 genome using STAR v2.6.1d. Transcript abundance was quantified using HTSeq v0.5.4p5.

Standard procedures for filtering, log-normalization, variable gene selection, dimensionality reduction and clustering were performed using the Seurat 3.0 package (*42*). Briefly, cells with fewer than 800 detected genes, more than 5,000 detected genes or more than 15% mitochondrial genes were discarded. To make counts comparable among cells, gene counts were normalized to 10,000 reads per cell, then log-transformed. Following PCA dimensionality reduction, cells were clustered by running the Louvain algorithm and visualized using UMAP. Differential expression analysis was performed using the Wilcoxon rank-sum test implemented in the Seurat package's 'FindAllMarkers' function. Significantly differentially expressed genes were defined as those with log fold change > 0.5 and Bonferroni-corrected p-value < 0.05.

**Single cell TCR-seq**

TCR-seq was performed using our previously developed single-cell paired TCR sequencing method (*45*) with some modifications. Briefly, for the first TCR reaction, 1 μL of the cDNA products of Smart-seq2 was preamplified with HotStarTaq DNA polymerase (Qiagen) using multiplex PCR with multiple Vα and Vβ region primers, Cα and Cβ region primers. A 25-cycle first PCR reaction was done per manufacturer's instructions using the following cycling conditions: 95 °C 15 min; 94 °C 30 s, 62 °C 1 min, 72 °C 1 min × 25 cycles; 72 °C 10 min; 4 °C. Next, 1 μL aliquot of the first reaction was used as a template for second 12 μL PCR using HotStarTaq DNA polymerase (Qiagen) with multiple internally nested TCRVα, TCRVβ, TCRCα and Cβ primers. The cycling conditions were: 95 °C 15 min; 94 °C 30 s, 64 °C 1 min, 72 °C 1 min × 25 cycles; 72 °C 7 min; 4 °C. 1 μL aliquot of the second PCR product was used as a template for the third 20 μL

PCR reaction, which incorporates barcodes and enables sequencing on the Illumina MiSeq platform. For the third and final PCR reaction for TCR sequencing, amplification was performed with HotStarTaq DNA polymerase for 36 cycles using a 5′ barcoding primer (0.05 μM) containing the common 23-base sequence and a 3′ barcoding primer (0.05 μM) containing sequence of a third internally nested Cα and/or Cβ primer, and Illumina Paired-End primers. The cycling conditions were: 95 °C 15 min; 94 °C 30 s, 66 °C 30 s, 72 °C 1 min × 36 cycles; 72 °C 10 min; 4 °C. The PCR products were combined at equal proportion by volume, run on a 1.2% agarose gel, and a band around 350 to 380 bp was excised and gel purified using a Qiaquick gel extraction kit (Qiagen). This purified product was then sequenced on a Miseq platform (Illumina) acquiring 2× 250 bp reads.

**Bulk TCRβ sequencing**

$KIR^+CD8^+$ T cells were sorted from PBMCs of 9 healthy subjects, and DNA was extracted using QIAamp DNA Micro Kit (Qiagen). Immunosequencing of the CDR3 regions of human TCRβ chains was performed using the immunoSEQ Assay by Adaptive Biotechnologies.

***In vitro* cell proliferation assay**

$CD8^+$ T cells were purified from PBMCs of healthy donors using CD8 microbeads (Miltenyi) per manufacturer's instructions, stained with flow antibodies, and live $CD3^+CD56^-CD8^+KIR^+$ or $KIR^-$ T cells were sorted out by FACSAria Fusion flow cytometer (BD). The sorted $KIR^+$ or $KIR^-$ $CD8^+$ T cells were stimulated with anti-CD3/CD28 beads (Gibco) at 1:1 ratio (1 μL beads per $4{\times}10^4$ cells) supplemented with 50 U/mL IL-2 in 96-well plates for 18 hours. The $CD8^-$ PBMCs were labeled with CellTrace Violet (CTV, ThermoFisher) per manufacturer’s instruction. 1 μg/mL anti-CD3 (UCHT-1) was coated on 96-well plate in 50 μL PBS per well at 4 °C overnight. After removal of anti-CD3/CD28 microbeads, $KIR^+$ and $KIR^-$ $CD8^+$ T cells were mixed with CTVlabelled $CD8^-$ PBMCs at 1:30 ratio and cultured in 96-well plate pre-coated with 1 μg/mL anti-CD3. After 4 or 6 days, cells were harvested and dilution of CTV in $CD4^+$ T cells was analyzed by flow cytometry.

**Statistical analysis.** No specific statistical methods were used to predetermine sample size. All results are presented as the mean ± SEM. The significance of the difference between groups was analyzed as described in the figure legends. Pearson's correlation coefficients with two-tailed *P* values were determined in the correlation analysis. $P$ values < 0.05 were considered statistically significant. All statistical analyses were performed using GraphPad Prism Software version 8.0.2.

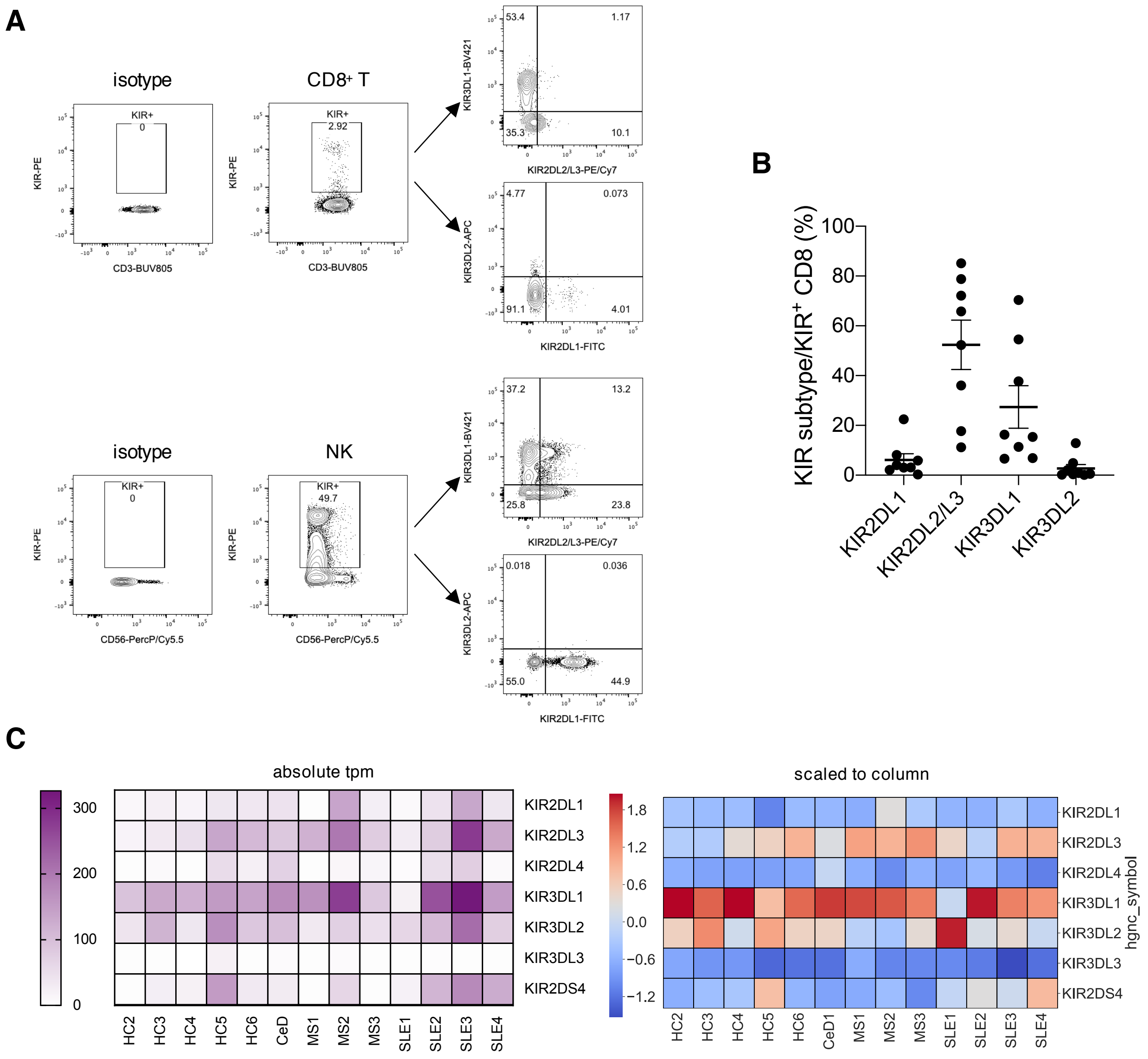


**Fig. S1. KIR3DL1 and KIR2DL3 are the two major KIR subtypes expressed on CD8$^+$ T cells.** **(A)** Representative plots showing expression of KIR subtypes among KIR$^+$CD8$^+$ T cells and KIR$^+$ NK cells in the peripheral blood, detected by flow cytometry. Total PBMCs were stained with PE-conjugated antibodies against KIR2DL1/S5, KIR2DL2/L3/S2, KIR2DL5, KIR3DL1, KIR3DL2 and FITC-conjugated anti-KIR2DL1/S5, PE/Cy7-conjugated anti-KIR2DL2/L3/S2, BV421-conjugated anti-KIR3DL1, APC-conjugated anti-KIR3DL2, together with other surface antibodies. (**B**) Summary scatter plot showing the percentage of KIR2DL1$^+$, KIR2DL2/L3$^+$, KIR3DL1$^+$ and KIR3DL2$^+$ cells among KIR$^+$CD8$^+$ T cells from the blood of healthy controls (N=8). (**C**) Transcript levels of different KIR genes in sorted KIR$^+$ CD8$^+$ T cells detected by RNA sequencing. Left panel shows absolute tpm (transcript per million reads) of the KIR genes in each individual, while right panel shows normalized expression level of the KIR genes (scaled to column) among different donors.

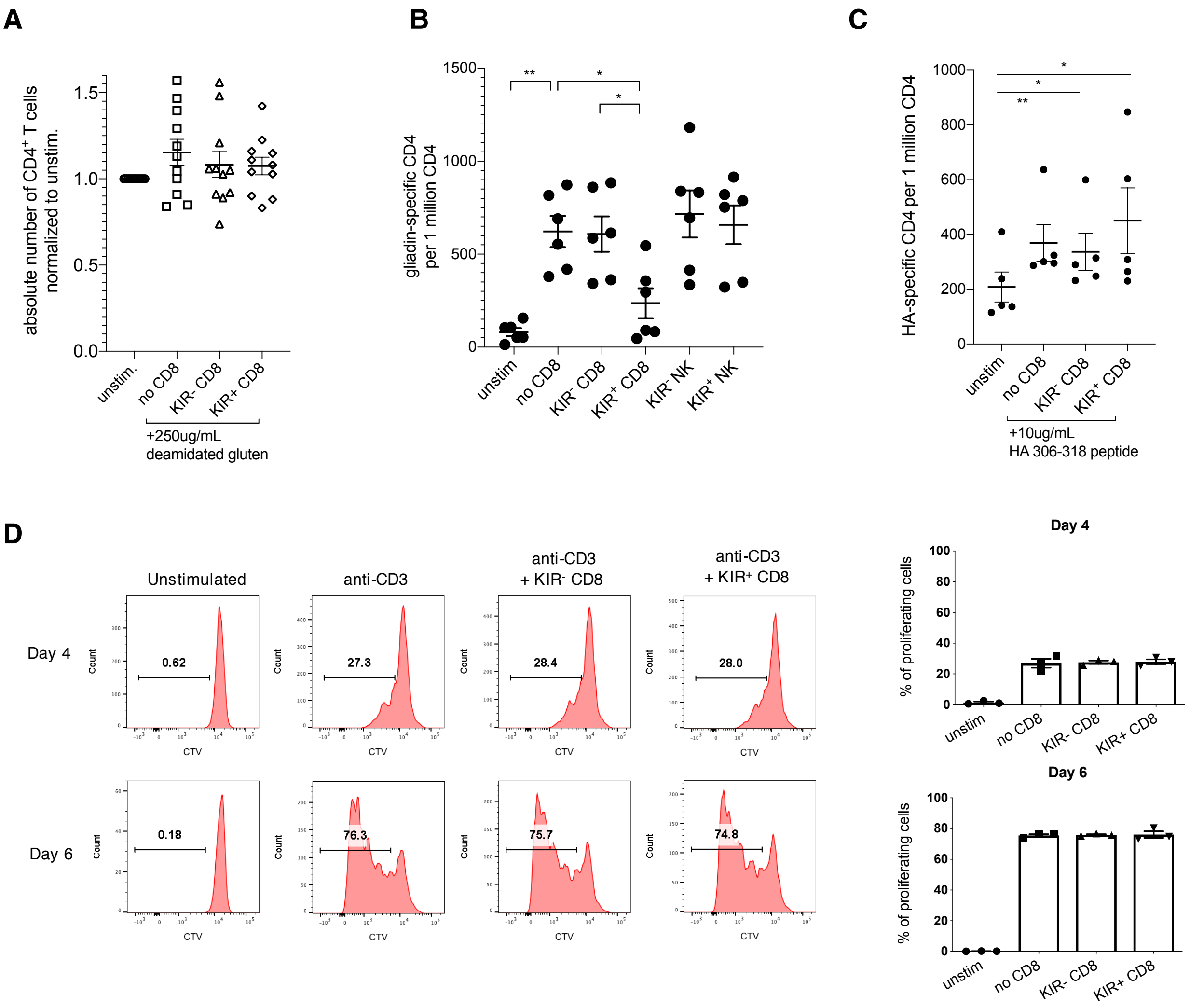


**Fig. S2. KIR⁺CD8⁺ T cells target gliadin-specific pathogenic CD4⁺ T cells specifically.** (**A**) Absolute number of total CD4⁺ T cells in the cultures of Day 6 (N=11) as described in Fig. 3A. (**B**) Frequency of gliadin-specific CD4⁺ T cells in the co-cultures of sorted KIR⁻CD8⁺ T cells, KIR⁺CD8⁺ T cells, KIR⁻ NK or KIR⁺ NK with CD8⁻ PBMCs from CeD patients (N=6) on Day 6 in responses to stimulation with 250ug/mL deamidated gluten. *$P$<0.05, **$P$<0.01, Friedman test corrected for multiple comparisons. (**C**) Frequency of HA-specific CD4⁺ T cells (detected by DRB1*04:Influenza A HA 306-318 tetramers) in the co-cultures of pre-activated KIR⁻ or KIR⁺ CD8⁺ T cells with CD8⁻ PBMCs from HLA-DR4⁺ healthy donors (N=5) on Day 6 in responses to stimulation with 10ug/mL HA 306-318 peptides. *$P$<0.05, **$P$<0.01, Friedman test corrected for multiple comparisons. (**D**) Representative figures and summary histograms showing proliferation of CD4⁺ T cells without stimulation or cultured with/without pre-activated KIR⁻ or KIR⁺ CD8⁺ T cells in the presence of plate-bound anti-CD3.

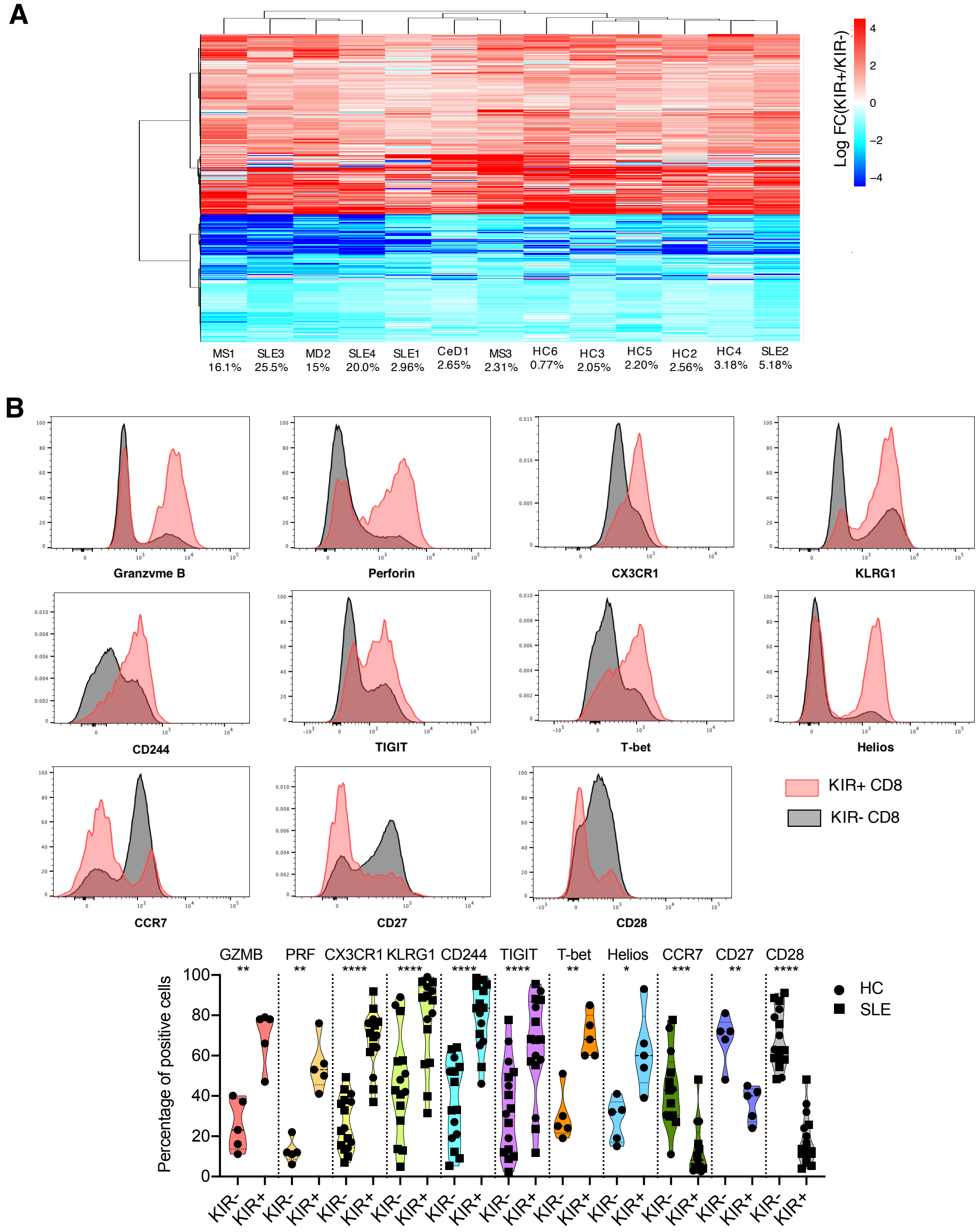


**Fig. S3. Phenotypic analysis of KIR$^+$ versus KIR$^-$ CD8$^+$ T cells in the blood.** (**A**) Heatmap showing fold changes of the 963 DEGs between KIR$^+$ and KIR$^-$ CD8$^+$ T cells from all individuals. Columns show samples annotated with frequency of KIR$^+$ CD8$^+$ T cells in total CD8$^+$ T cells, and rows and columns are ordered based on hierarchical clustering. Normalized fold changes are centered for each gene. (**B**) Protein expression of featured DEGs (Granzyme B (GZMB), perforin (PRF), CX3CR1, KLRG1, CD244, TIGIT, T-bet, Helios, CCR7, CD27 and CD28) in KIR$^+$ and KIR$^-$ CD8$^+$ T cells from the blood of healthy controls (HC) and patients with systemic lupus erythematosus (SLE) measured by flow cytometry. **P*<0.05, ***P*<0.01, ****P*<0.001, *****P*<0.0001, mixed-effects analysis followed by Holm-Sidak's multiple comparisons test.

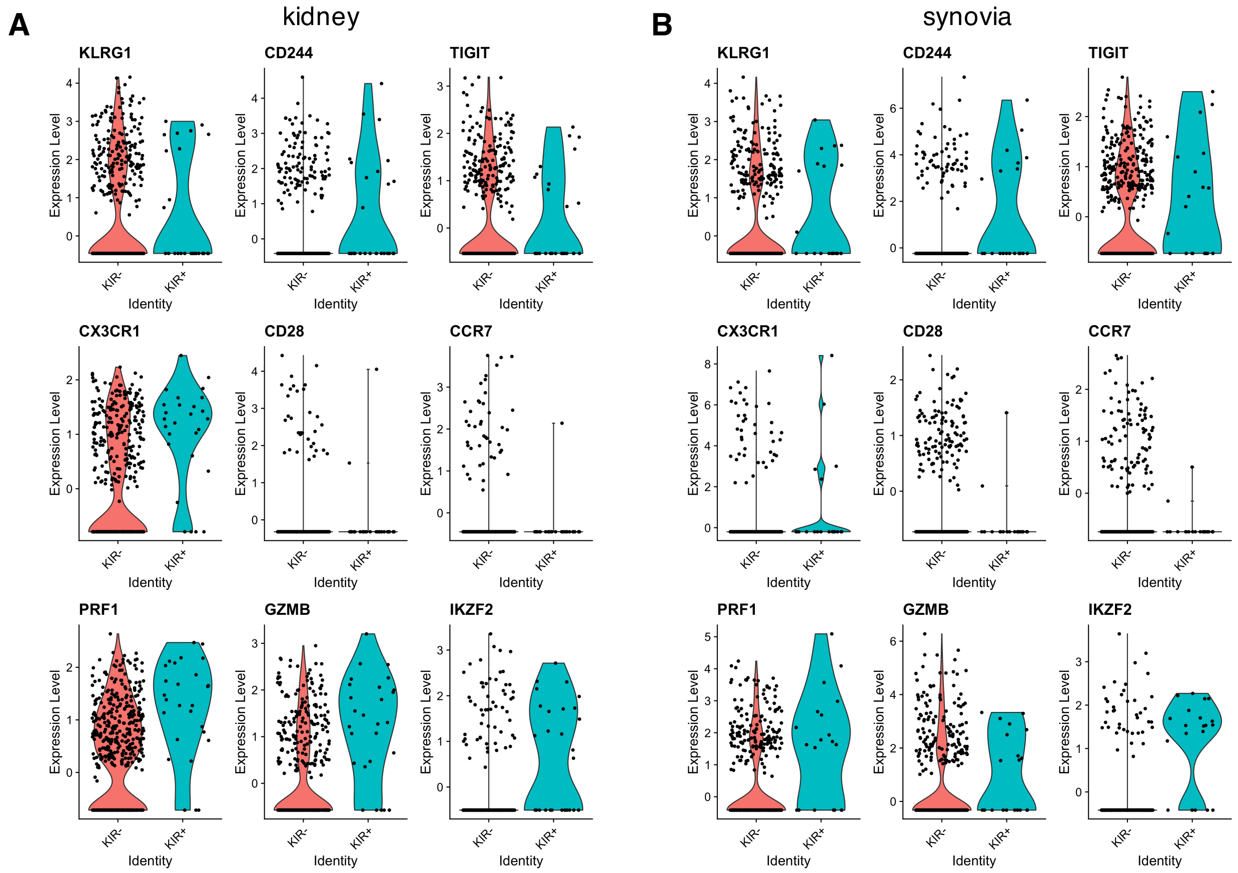


**Fig. S4. Phenotypic analysis of KIR$^+$CD8$^+$ T cells in inflamed tissues.** KIR$^+$ versus KIR$^-$ CD8$^+$ T cells in the kidney of SLE patients or synovial tissues from rheumatoid arthritis (RA) were analyzed using the single cell RNA-seq data generated by AMP RA/SLE program. The UMI count matrix was imported and CD8$^+$ T cells (expressing *CD3E*, *CD8A* and *CD8B* transcripts) were selected for further analysis. Violin plots showing expression of *KLRG1*, *CD244*, *TIGIT*, *CX3CR1*, *CD28*, CCR7, *PRF1*, *GZMB* and *IKZF2* transcripts in KIR$^+$ (expressing any of the *KIR* transcripts: *KIR3DL1*, *KIR2DL3*, *KIR2DL2*, *KIR2DL1* or *KIR3DL2*) versus KIR$^-$ CD8$^+$ T cells in the SLE kidney (**A**) and RA synovia (**B**).

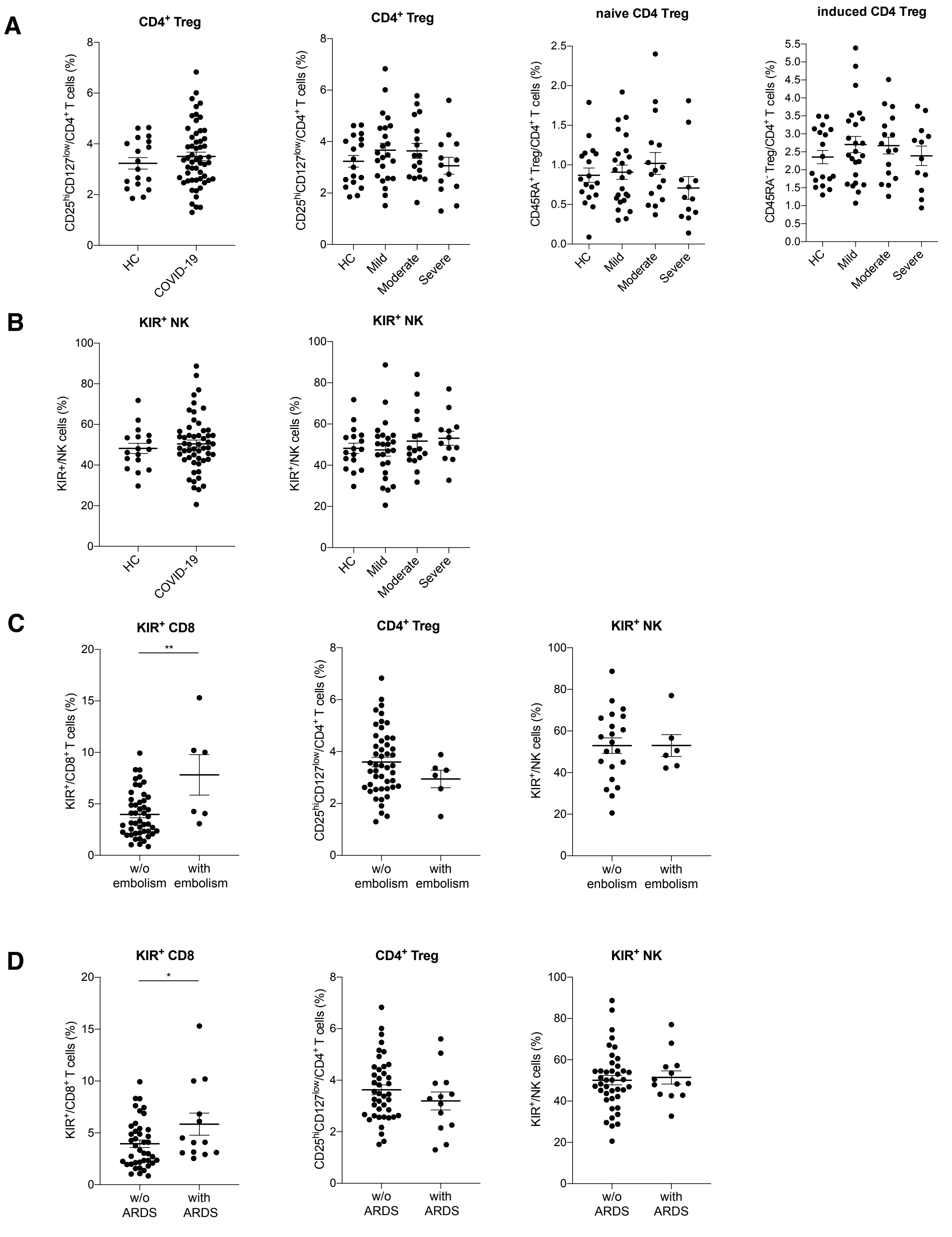
A
CD4+ Treg
CD25hiCD127low/CD4+ T cells (%)
HC
COVID-19
Mild
Moderate
Severe
naive CD4 Treg
CD45RA+ Treg/CD4+ T cells (%)
induced CD4 Treg
CD45RA- Treg/CD4+ T cells (%)
B
KIR+ NK
KIR+/NK cells (%)
C
KIR+ CD8
KIR+/CD8+ T cells (%)
**
w/o embolism
with embolism
w/o enbolism
D
*
w/o ARDS
with ARDS

**Fig. S5. Flow cytometric analysis of PBMCs from COVID-19 patients.** (**A**) Summary histograms showing frequency of $CD4^+$ regulatory T cells ($CD25^{hi}CD127^{low}$) as well as subpopulations of $CD4^+$ Tregs in the peripheral blood of healthy controls (HC, N=17) and COVID-19 patients with varying disease severity (Mild: 23, Moderate: 17, Severe: 13). (**B**) Summary histograms showing frequency of $KIR^+$ NK cells in the peripheral blood of healthy controls (HC, N=17) and COVID-19 patients with varying disease severity (Mild: 23, Moderate: 17, Severe: 13). (**C**) Frequency of $KIR^+CD8^+$ T cells, $CD4^+$ Tregs ($CD25^{hi}CD127^{low}$) and $KIR^+$ NK cells in COVID-19 patients with or without embolism. (**D**) Frequency of $KIR^+CD8^+$ T cells, $CD4^+$ Tregs ($CD25^{hi}CD127^{low}$) and $KIR^+$ NK cells in COVID-19 patients with or without ARDS.

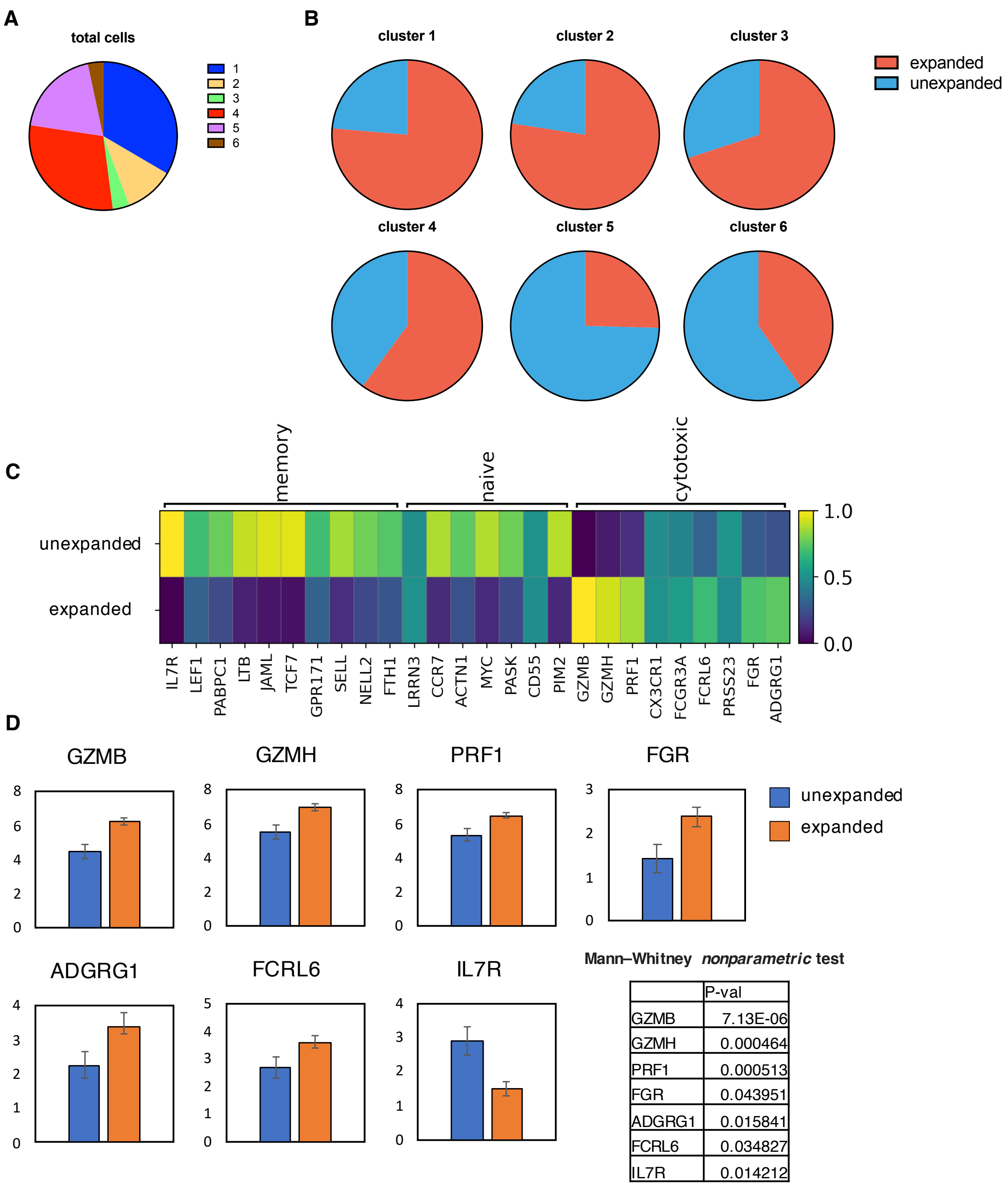


| | P-val |
|---|---|
| GZMB | 7.13E-06 |
| GZMH | 0.000464 |
| PRF1 | 0.000513 |
| FGR | 0.043951 |
| ADGRG1 | 0.015841 |
| FCRL6 | 0.034827 |
| IL7R | 0.014212 |

**Fig. S6. Single cell RNA-seq analysis of KIR⁺CD8⁺ T cells.** (**A**) Pie chart depicting the composition of the 6 clusters identified by unsupervised clustering in total KIR⁺CD8⁺ T cells. (**B**) Pie charts showing the composition of expanded and unexpanded KIR⁺CD8⁺ T cells in each cluster. **C**-**D**, Single-cell RNA-seq analysis of expanded versus unexpanded KIR⁺CD8⁺ T cells from COVID-19 patients by 10x Genomics. (**C**) Heatmap displaying normalized expression of naïve-, memory- or cytotoxic-associated genes in unexpanded versus expanded KIR⁺CD8⁺ T cells from the blood of COVID-19 patients. (**D**) Bar graphs showing mRNA expression of *GZMB*, *GZMH*, *PRF1*, *FGR, ADGRG1, FCRL6* and *IL7R* in unexpanded and expanded KIR⁺CD8⁺ T cells in the peripheral blood from COVID-19 patients.

**Table S1. (separate file)**
Differentially expressed genes between $KIR^+$ and $KIR^-$ CD8+ T cells from MS patients

**Table S2. (separate file)**
Differentially expressed genes between $KIR^+$ and $KIR^-$ CD8+ T cells from healthy donors and a variety of autoimmune diseases (MS, SLE and CeD)

**Table S3. (separate file)**
Differentially expressed features of each cluster compared to all other cells among $CD8^+$ T cells from healthy controls, MS patients and COVID-19 patients (10x Genomic platform)

**Table S4. (separate file)**
Differentially expressed features of each cluster compared to all other cells among $KIR^+CD8^+$ T cells from healthy subjects and patients with autoimmune diseases (Smart-seq2 platform)

**Table S5. (separate file)**
Detailed information of patients and healthy controls included in the study

**Table S6. (separate file)**
Clinical metadata of COVID-19 patients and healthy controls included in the study
